# Necrotizing enterocolitis is preceded by increased gut bacterial replication, *Klebsiella*, and fimbriae-encoding bacteria that may stimulate TLR4 receptors

**DOI:** 10.1101/558676

**Authors:** Matthew R. Olm, Nicholas Bhattacharya, Alexander Crits-Christoph, Brian A. Firek, Robyn Baker, Yun S. Song, Michael J. Morowitz, Jillian F. Banfield

**Affiliations:** Department of Plant and Microbial Biology, University of California, Berkeley, CA, USA; Department of Mathematics, University of California, Berkeley, CA, USA; Department of Surgery, University of Pittsburgh School of Medicine, Pittsburgh, PA, USA; Division of Newborn Medicine, UPMC Magee-Womens Hospital, Pittsburgh, PA, USA; Department of Statistics, University of California, Berkeley, CA, USA; Department of Electrical Engineering and Computer Sciences, University of California, Berkeley, CA, USA; Department of Earth and Planetary Science, University of California, Berkeley, CA, USA; Department of Environmental Science, Policy, and Management, University of California, Berkeley, CA, USA; Earth Sciences Division, Lawrence Berkeley National Laboratory, Berkeley, CA, USA; Chan Zuckerberg Biohub, San Francisco, CA, USA

## Abstract

Necrotizing enterocolitis (NEC) is a devastating intestinal disease that occurs primarily in premature infants. We performed genome-resolved metagenomic analysis of 1,163 fecal samples from premature infants to identify microbial features predictive of NEC. Features considered include genes, bacterial strain types, eukaryotes, bacteriophages, plasmids and growth rates. A machine learning classifier found that samples collected prior to NEC diagnosis harbored significantly more *Klebsiella*, bacteria encoding fimbriae, and bacteria encoding secondary metabolite gene clusters related to quorum sensing and bacteriocin production. Notably, replication rates of all bacteria, especially Enterobacteriaceae, were significantly higher two days before NEC diagnosis. The findings uncover biomarkers that could lead to early detection of NEC and targets for microbiome-based therapeutics.

## Main

Necrotizing enterocolitis (NEC) is widely studied yet poorly understood. First described in the early 1800s^1^, NEC is a disorder of intestinal inflammation that can progress to bowel necrosis, sepsis, and death^2^. NEC affects 7% of very low birthweight infants born in the United States each year, and mortality rates have remained around 20 – 30% for several decades^2^. The direct cause or causes of NEC remain unknown.

The primary risk factor for NEC is preterm birth^2^. Immature enterocytes exhibit hyperactive immune responses through the TLR4 pathway in response to bacterial lipopolysaccharide (LPS)^3^, which can lead to bowel damage^4^. Experimental NEC occurs in conventionally raised animals but not those reared in a germ-free environment^5,6^. These observations suggest that the intestinal microbiome plays a role in the disease and led to the prevailing hypothesis that an excessive immune response to abnormal gut microbes is the most likely basis for the pathogenesis of NEC. Although no single microbe has been consistently identified as a biomarker for NEC^7^, increased abundance of bacteria in the phylum Proteobacteria is a frequently reported microbial pattern in NEC infants^8^. Most fecal microbiome-based profiling studies of NEC utilize 16S rRNA amplicon sequencing, which provides a general overview of the bacteria present, but does not reveal metabolic features that could contribute to NEC pathogenesis.

Genome-resolved methods may provide new insights into NEC development. The approach has several advantages over 16S rRNA amplicon sequencing. All DNA in a sample is sequenced, allowing detection of bacteriophages, plasmids, eukaryotes, and viruses. Bioinformatics techniques can also infer *in situ* bacterial replication rates directly from metagenomic data^9,10^, an important metric, as some microbiome-related diseases have a signal related to bacterial replication but not relative abundance^10^. Importantly, genome assembly and annotation can provide functional information about organisms present and possibly reveal genes associated with NEC. Further, whole-genome comparisons provide strain discrimination, and thus detailed testing of Koch’s postulates. Finally, mapping to reference genomes is not required for genome detection, allowing for the discovery of novel bacterial clades^11^. While identification of a single causative strain, virus, or toxin would be the most actionable result for clinicians, any associations could potentially be used as biomarkers to identify early warning signs of NEC, and microbial communities associated with NEC could be targeted with microbiome-altering techniques such as probiotics, prebiotics, or other approaches^12^.

## Results

### Metagenomic characterization of premature infant fecal samples

We analyzed 1,163 fecal metagenomes from 34 preterm infants that developed NEC and 126 preterm infants without NEC (Figure 1). Premature infant subjects were matched for gestational age and calendar date, and recruited from the UPMC Magee-Womens Hospital (Pittsburgh, PA) over a 5 year period. Fecal samples were banked and specific samples were later chosen for DNA extraction and sequencing to preferentially study samples immediately prior to NEC. An average of 7.2 samples per infant, mostly from the first month of life, were sequenced and a total of 4.6 tera base pairs of shotgun metagenomic sequencing generated (**Supplemental Table S1**). Detailed sequencing information (**Supplemental Table S1**) and patient metadata (**Supplemental Table S2**) are provided.

**Figure 1:**
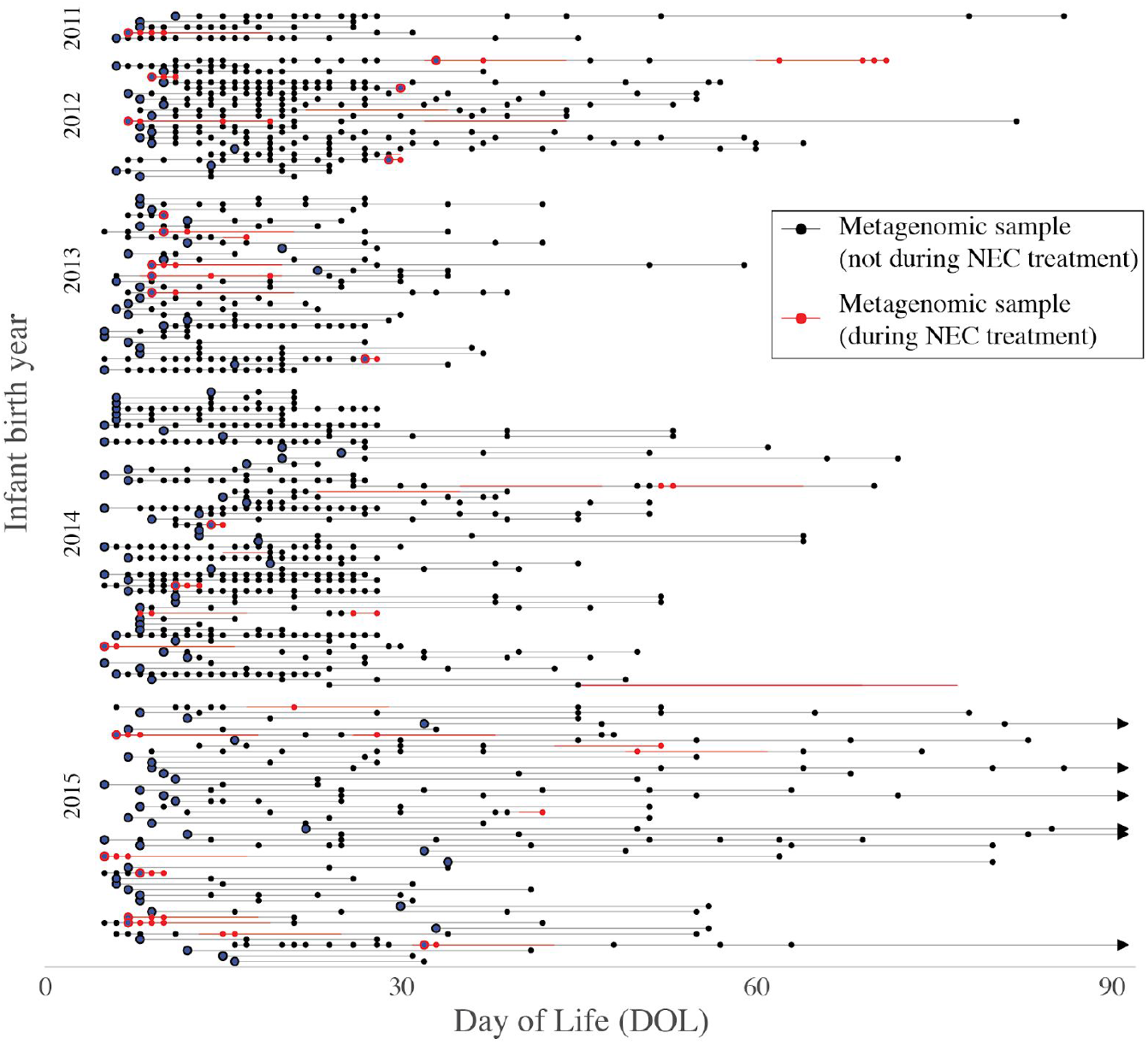
Metagenomic characterization of 1,163 samples from 160 premature infants. Each infant is represented by a horizontal line, and dots on the line represent sequenced metagenomic samples. Red sections indicate periods in which the infant was undergoing treatment for necrotizing enterocolitis. For some statistical tests one sample was chosen for each infant (pre-NEC and control samples); these samples are marked with larger circles.

Extensive computational analyses were performed on all samples to recover genomes *de novo*, and determine their phylogeny, metabolic potential, and replication rates (iRep^9^). We also searched samples for eukaryotic viruses, virulence factors, secondary metabolite gene clusters, and previously implicated pathogens^13,14^ (Figure 1a). This analysis resulted in 36 gigabase pairs of assembled sequence, 2,425 de-replicated bacterial genomes (average of 92% completeness and 1. 1% contamination), 5,218 bacteriophage genomes, 1,183 plasmid genomes, 7 eukaryotic genomes, and 804,185 *de novo* protein clusters (Figure 1b**; Supplemental Table S6**). As NEC can be a rapidly progressive disorder, for most statistical tests we defined NEC samples as those taken within 2 days prior to NEC diagnosis (“pre-NEC” samples). For infants that did not develop NEC, only one sample from the period associated with NEC onset was used (“control” samples). Pre-NEC and control samples were matched for day of life (DOL), gestational age, and recent antibiotic administration (Figure 1c**; Supplemental Figure S1**). For other analyses, when explicitly stated, all samples were used.

### Klebsiella pneumoniae is enriched in samples from infants with NEC

The gut microbiomes of all infants were dominated by Proteobacteria, regardless of NEC development (Figure 2a,b). As compared to previous studies of full-term infants^15,16^, the premature infants in this study had increased Enterobacteriaceae (a family of Proteobacteria to which many nosocomial pathogens belong^17^) and notably low abundances of Actinobacteria and Bacteroidetes. Factors that could select for these organisms include prophylactic antibiotics given to all premature infants at birth, high rates of birth by cesarean-section, predominance of formula feeding and immaturity of the intestine and immune system. Only the NEC microbiomes contained Fusobacteria and Tenericutes (Figure 2). Compared to control infants, the NEC infant microbiomes exhibited less stability, lower levels of Firmicutes, and higher levels of Enterobacteriaceae than the microbiomes in control infants (*p* = 8.9E-7; Wilcoxon rank sums test; **Supplemental Table S9;** Figure 2a). The general association of Enterobacteriaceae and infants that go on to develop NEC has been described previously^18^, but this prior analysis was not restricted to the period immediately prior to NEC detection. In our study, the gut microbiomes of infants that developed NEC were not significantly enriched in Enterobacteriaceae in pre-NEC vs. control samples (*p* = 0.15; Wilcoxon rank sums test), so the association of Enterobacteriaceae and NEC infants overall may be due to the proliferation of these bacteria after administration of antibiotics to treat NEC (**Supplemental Figure S9**).

**Figure 2:**
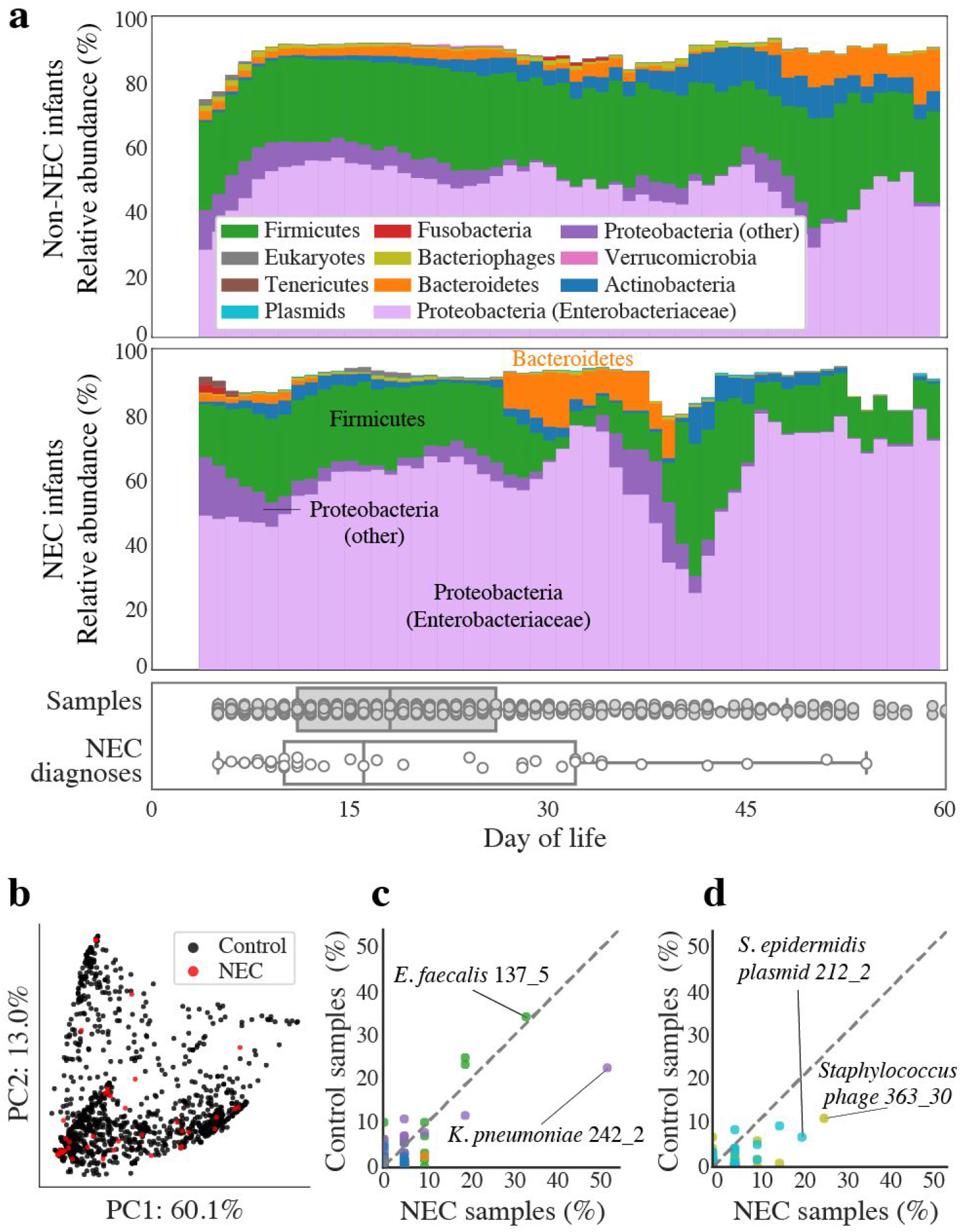
Comparison of microbes in premature infants that do and do not develop NEC. **(a)** The compositional profile of microbes colonizing infants that were and were not diagnosed with NEC. Bacteria were classified based on their phyla and other microbes were classified based on their domain. Each color represents the percentage of reads mapping to all organisms belonging to a taxon, and the stacked boxes for each sample show the fraction of reads in that dataset accounted for by the genomes assembled from the sample. Proteobacteria were subdivided into the family Enterobacteriaceae and other. All relative abundance values were averaged over a 5-day sliding window. Boxplots show the days of life in which samples were collected (top) and in which infants were diagnosed with NEC (bottom). **(b)** Principal component analysis (PCA) based on weighted UniFrac distance for all samples from NEC infants (red) and control infants (black). **(c, d)** The percentage of NEC infants vs. the percentage of non-NEC infants colonized by strains of **(c)** bacteria or **(d)** bacteriophage (gold) and plasmids (blue). Colonization by bacteria is defined as the presence of a strain at ≥ 0.1% relative abundance. Plasmid and bacteriophage detection required read-based genome breadth of coverage of ≥50%. Each dot represents a strain, and dashed lines show a 1:1 colonization rate.

A principal component analysis based on weighted UniFrac distance was performed to compare the microbiomes of all samples from all time points (Figure 2b). The first two principal components explained 73% of the overall variance, but samples collected from NEC infants (red) did not cluster separately from control infants (black dots). Consideration of higher principal components (up to the 5^th^ principal component) did not separate pre-NEC and control samples, and samples coded by clinical metadata also did not cluster together (**Supplemental Figure S7**).

To identify strains enriched in pre-NEC samples, the percentage of pre-NEC vs. control samples carrying each assembled bacterial, bacteriophage, and plasmid genome was calculated (Figure 2c,d). *Klebsiella pneumoniae* strain 242_2 was the most associated with NEC, and was present above the threshold of detection in 52% of pre-NEC samples vs. 23% of control samples (*p* = 0.008; Fisher’s exact test) (**Supplemental Table S12**). Interestingly, closely related bacteria (>99% average nucleotide identity (ANI)) colonized up to 35% of all infants (Figure 2c). This is likely the result of colonization of multiple infants by the same hospital-associated bacteria^19^. Importantly, no organisms in this study satisfied Koch’s postulate that a disease causing organism should be found in all NEC infants and no healthy patients.

### Bacterial replication rates are higher prior to NEC development

Bacterial replication rates are measured from metagenomic data by determining the difference in DNA sequencing coverage at the origin vs. terminus of replication, yielding an index of replication (iRep) that correlates with traditional doubling time measurements^9,10^. Remarkably, iRep values of bacteria overall were significantly higher in pre-NEC vs. control samples (*p* = 0.0003; Wilcoxon rank sums test), in a cohort balanced for DOL, gestational age, and recent antibiotic administration (Figure 3) (**Supplemental Table S9**). Further, iRep values followed a striking pattern in relation to NEC diagnosis: bacterial replication was stable four or more days prior to NEC diagnosis, increased daily in the three days prior to diagnosis, and crashed following diagnosis (probably due to resulting antibiotic administration) (Figure 3a). Individual species did not have enough data-points to be plotted confidently (minimum of 5 measurements per DOL), but genomes of the family Enterobacteriaceae displayed similar but more dramatic patterns than other bacteria overall (Figure 3ab). Increased bacterial replication prior to NEC could promote disease onset or merely be a reaction to changing conditions in the gut that led to NEC.

**Figure 3.**
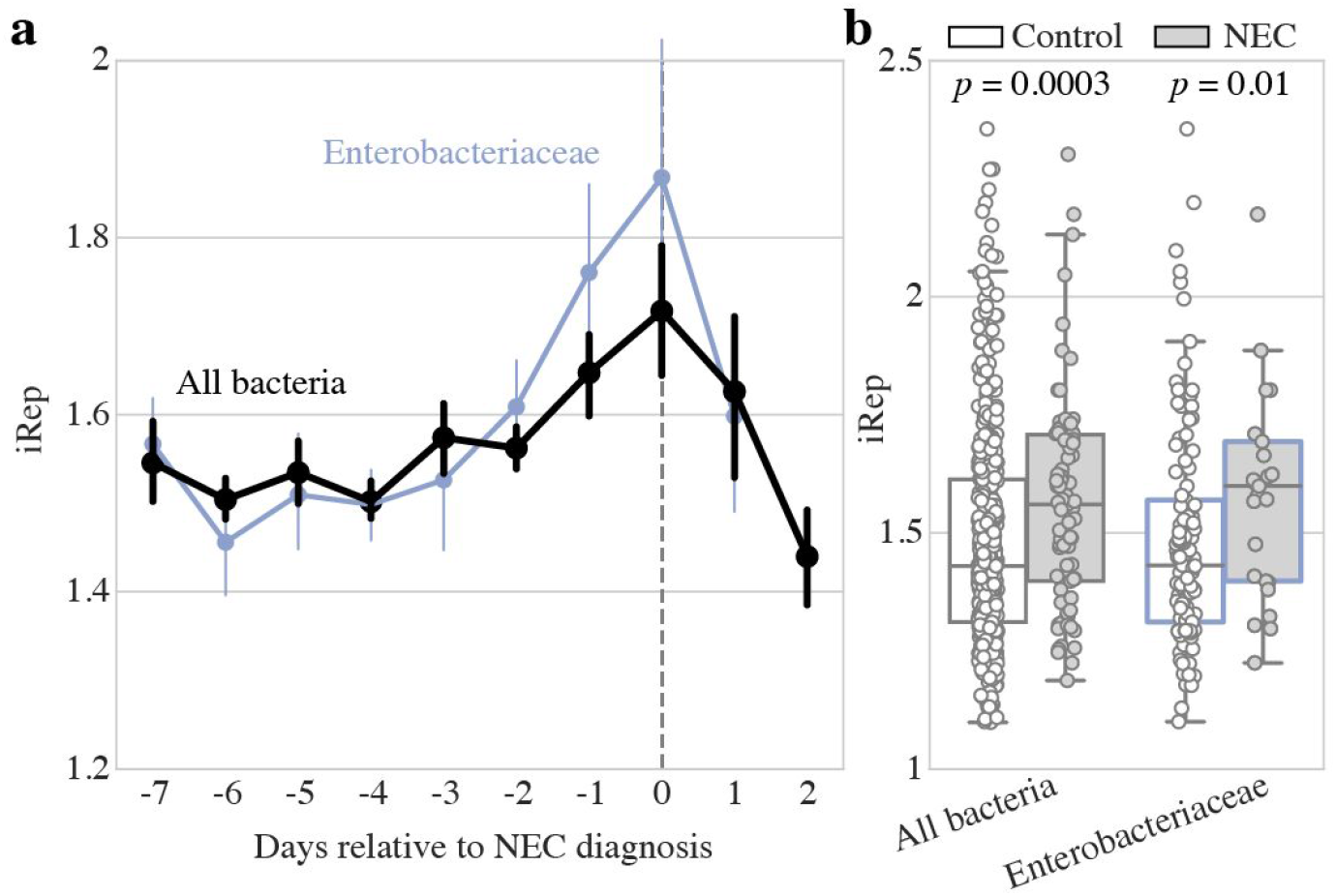
Bacterial replication rates are significantly higher prior to NEC development. **(a)** Replication rates for bacterial groups relative to day of NEC diagnosis. Dots represent the mean value for each group on each day, and error bars represent standard error of the mean. Days of life in which growth rates were calculated from at least 5 infants are shown. **(b)** Growth rates in control (white) vs pre-NEC (grey) samples. *p*-values shown from Mann–Whitney U test.

### Machine learning identifies additional differences between NEC and control cases

2,119 features (e.g., the abundance of organisms encoding secondary metabolite gene clusters and each of 600 KEGG modules (specific metabolic pathways)) were measured for each of the 1,163 metagenomic samples (Figure 1**; Supplemental Table S3**). In order to evaluate which features are most different between pre-NEC and control samples, a machine learning (ML) classifier was developed. Multiple ML algorithms were evaluated, and although all performed with similar accuracy (**Supplemental Table S10**), the boosted gradient classifier was ultimately chosen due to its known ability to handle class imbalance. The classifier was trained on all 2,119 features to predict if samples were pre-NEC or control, and accuracy was measured through cross-validation over 100 iterations. The classifier achieved a median accuracy of 64% on balanced sets; 14% better than random chance. While a classifier with this accuracy may have limited utility in a clinical setting, it allowed us to interrogate which features were most informative for differentiating pre-NEC and control samples.

The most important individual features used by the ML classifier were replication rates (iRep values), KEGG modules, secondary metabolite gene clusters, and overall plasmid abundance (Figure 4). iRep values of both specific bacterial taxa and median iRep values overall were some of the most important features (Figure 4b), while KEGG modules accounted for over 50% of the total feature importance (Figure 4a) (**Supplemental Table S4**). A similar number of KEGG modules were associated and anti-associated with NEC (Figure 4c), but descriptions of the pathways associated with NEC (e.g., erythritol and galactitol transport systems) and anti-associated with NEC (e.g., sodium and capsular polysaccharide transport systems) bear no obvious relationship to the disease (**Supplemental Table S4**). Secondary metabolite gene clusters were the second most important category overall (Figure 4a), but unlike KEGG modules, very few were anti-associated with NEC (Figure 4c). The most significant secondary metabolite gene cluster encodes an unusual operon of biosynthetic genes found in *Klebsiella* (cluster 416). In other species, similar operons are implicated in biosynthesis of quorum sensing butyrolactones^20^. The second most significant cluster of genes occurs in *Enterococcus* and is involved in biosynthesis of a sactipeptide resembling subtilosin A1, an antimicrobial agent with known hemolytic activity^21^ (cluster 438) (**Supplemental Table S5**). Interestingly, another cryptic secondary metabolite gene cluster with a high feature importance (cluster 432) is closely related to a previously characterized cluster on a plasmid of Enterotoxin-producing *Clostridium perfringens* adjacent to the enterotoxin gene (cpe) and beta2 toxin gene (cpb2)^22^. Overall, high plasmid abundance was correlated with pre-NEC samples (Figure 4b) and *K. pneumoniae* plasmids in particular were significantly more abundant in pre-NEC samples (*p* = 0.03) (**Supplemental Figure S3**).

**Figure 4:**
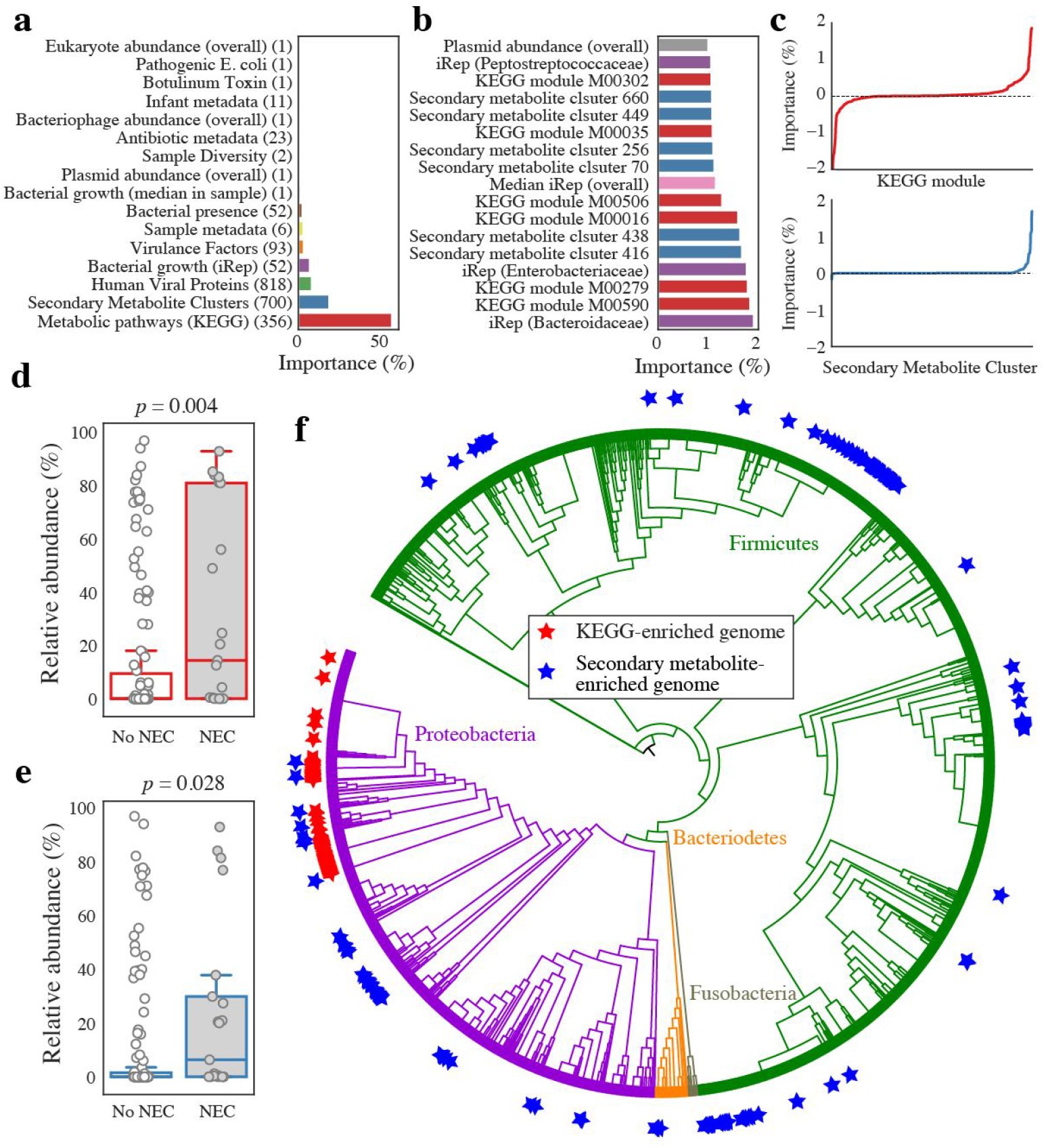
Machine learning identifies differences between pre-NEC and control samples. **(a)** The sum of all individual importances for each feature category. The number of features in each category is listed in parentheses. **(b)** Importance of all individual features associated with NEC with classifier importances over 1%. **(c)** Signed importances of all individual KEGG modules (top, red) and secondary metabolite clusters (bottom, blue). Negative values are negatively associated with pre-NEC samples, positive values are positively associated with pre-NEC samples. **(d, e)** The relative abundance of genomes enriched in important KEGG modules **(d)** and important secondary metabolite enriched genomes **(e)** in pre-NEC vs. control samples. *p*-values shown from Mann–Whitney U test. **(f)** The distribution of genomes enriched in important KEGG modules (red star) and important secondary metabolite clusters (blue star) around a phylogenetic tree of all recovered bacterial genomes. Genomes enriched in important KEGG modules are more clustered on the tree than those enriched in important secondary metabolite clusters.

Feature importances were also analyzed in combination. Each bacterial strain was assigned an importance value based on the sum of the importance scores for the KEGG modules encoded by its genome. 150 genomes have high KEGG importance values (hereinafter referred to as “organisms of interest”) (**Supplemental Table S7; Supplemental Figure S4**). Interestingly, the organisms of interest were significantly more abundant in pre-NEC samples as compared to control samples (*p* = 0.004) (Figure 4d), and they cluster phylogenetically (Figure 4f). 97% were in the family Enterobacteriaceae, and of those, 90% were in the genus *Klebsiella.* The prevalence of *K. pneumoniae* in pre-NEC samples (Figure 2c) may explain the high abundance of *K. pneumoniae* plasmids in these samples.

Secondary metabolite biosynthetic gene clusters identified to be important by the ML classifier occur in 218 organisms that are significantly associated with pre-NEC samples (Figure 4e). Several types of secondary metabolite gene clusters were enriched in this these genomes (p < 0.01; Fisher’s exact test), including sactipeptides, bacteriocins, and butyrolactones (encoded by 382, 286, and 11 genomes, respectively) (**Supplemental Table S7**). As opposed to organisms of interest, these bacteria were spread around the phylogenetic tree (Figure 4f). This may indicate that the clusters themselves are associated with pre-NEC samples. Overall, the results point to quorum sensing and anti-microbial peptide production as being associated with NEC onset.

### Bacteria associated with NEC encode specific types of fimbriae

We leveraged the gene content information provided by genome-resolved metagenomics to search for proteins associated with 1) pre-NEC samples and 2) organisms of interest. Three clustering algorithms were evaluated for their ability to reconstruct known clusters of ribosomal proteins (**Supplemental Table S11**), and a hybrid Markov Cluster algorithm approach^23^ performed best. Application of the algorithm to the 36,701,491 proteins reconstructed in this study yielded 804,277 protein clusters, none of which was statistically associated with NEC (Fisher’s exact test with false discovery rate correction) (Figure 5a). However, 85 protein clusters were associated with organisms of interest with high precision and recall (>0.7) (Figure 5b). The most common pFam annotations for these clusters were fimbriae and ABC transport proteins (**Supplemental Table S8**). However, only genomes encoding fimbrial proteins also had a significant association with NEC (*p* = 0.02; Wilcoxon rank-sum with Benjamini-Hochberg FDR correction; **Supplemental Table S8**).

**Figure 5.**
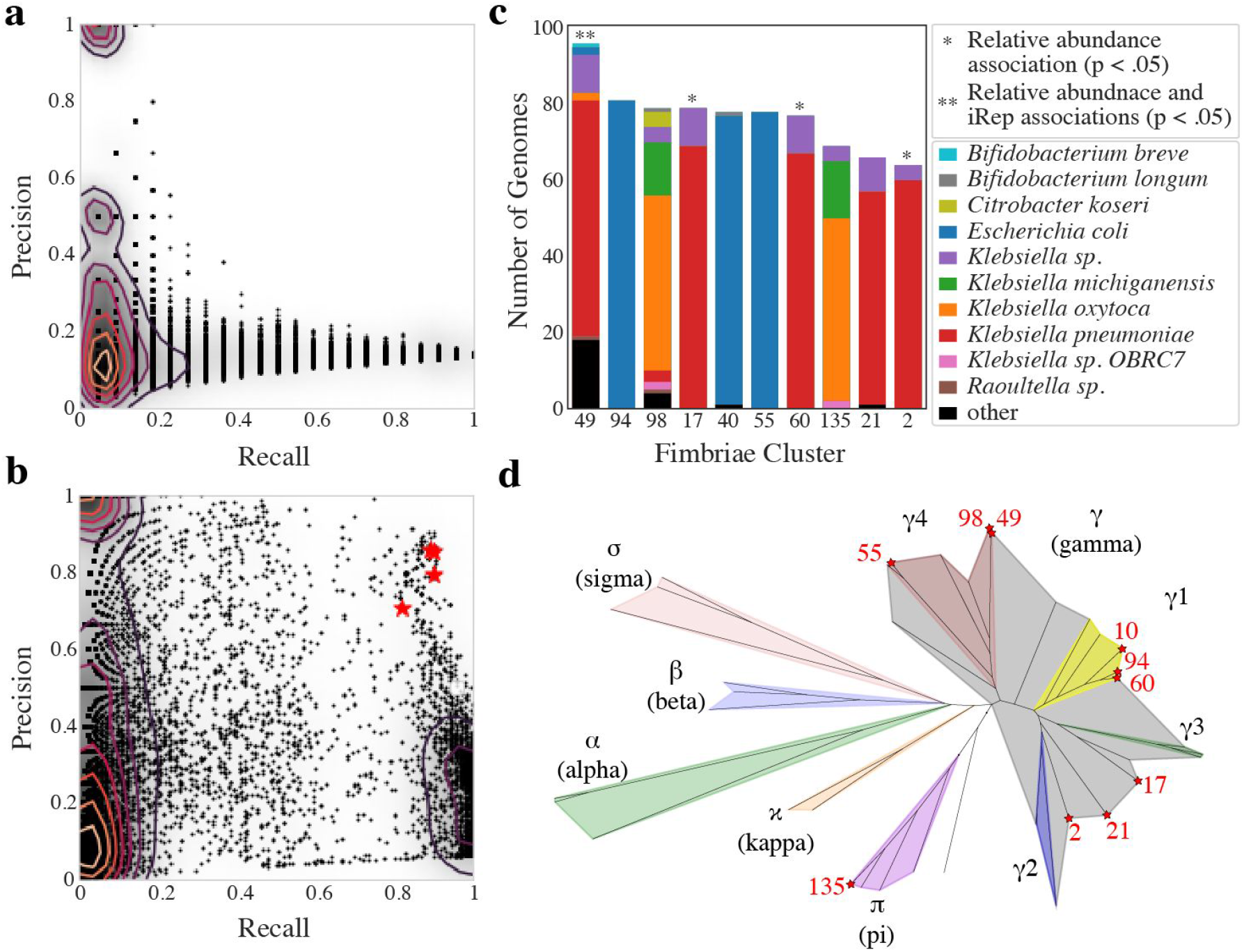
Genomes encoding fimbriae are associated with necrotizing enterocolitis development. **(a, b)** Association of protein clusters with pre-NEC samples **(a)** and organisms of interest **(b)**. Each dot represents a protein cluster. Recall is **(a)** the number of pre-NEC samples the cluster in / the total number of pre-NEC samples and **(b)** the number of organisms of interest the cluster in / the total number of organisms of interest. Precision is **(a)** the number of pre-NEC samples the cluster is in / the number of total pre-NEC samples, and **(b)** the number of organisms of interest the cluster is in / the total number of genomes the cluster is in. Clusters annotated as fimbriae are marked with a red star. Contour lines are drawn to indicate density. **(c)** The number of bacterial genomes encoding each fimbriae cluster, the special-level phylogenetic profile of genomes encoded by each fimbriae cluster, and each cluster’s association with NEC. **(d)** Phylogenetic tree of CU usher proteins built using IQtree. Three amino acid sequences from each *de novo* CU cluster and three reference amino acid sequences from each defined CU clade was included in the tree. Colors mark the phylogenetic breadth spanned by reference sequences, and stars represent *de novo* CU clades. For all *de novo* clusters the three randomly chosen sequences fell extremely close to each other on the tree.

Comparison of fimbrial operons against public databases revealed that the majority encode chaperone-usher (CU) type fimbriae. A classification scheme exists for CU fimbriae based on usher protein pFam PF00577.19^24,25^. The 32,646 usher proteins identified in our sequencing data (**Supplemental Table S13**) were clustered into groups based on amino acid sequence identity, and the ten most prevalent groups were placed in a phylogenetic tree with reference sequences from each subtype of CU fimbriae (Figure 5d). All ten fimbriae clusters fit into the established CU fimbriae taxonomy, with 9/10 falling in the γ super-clade and one into the π clade (Figure 5d). Four fimbriae clusters identified in this study were significantly more abundant in pre-NEC samples, and genomes encoding cluster 49 (γ4 clade) also had significantly higher iRep values in pre-NEC samples (Figure 5c). 27 genomes that encode fimbrial cluster 49 were not identified as genomes of interest, yet they were at significantly higher abundance, and have significantly higher iRep values, when considering all samples from NEC vs. control infants (**Supplemental Figure S6**) (*p* < 0.01; Wilcoxon rank-sums test). This suggests fimbrial cluster 49 itself may be associated with NEC and not incidentally associated with metabolically important genomes.

### Biomarkers of NEC are most informative closer to NEC diagnosis

Statistical tests uncovered four factors significantly associated with pre-NEC samples (samples taken within day days prior to NEC diagnosis): iRep values overall (Figure 3b), genomes encoding specific types of secondary metabolite gene clusters (sactipeptides, bacteriocins, and butyrolactones) (**Supplemental Table S7**), *Klebsiella* (Figure 2c), and fimbriae cluster 49 (Figure 5c). We performed a similar analysis each day up to eight days prior to NEC diagnosis (Figure 6). Genomes encoding specific types of secondary metabolite gene clusters and *Klebsiella* genomes were always significantly more abundant in NEC samples, though the effect size of the difference became slightly higher closer to NEC diagnosis. iRep values and the abundance of genomes encoding fimbriae cluster 49, on the other hand, were only significantly higher 3 days and 1 day prior to diagnosis, respectively.

**Figure 6.**
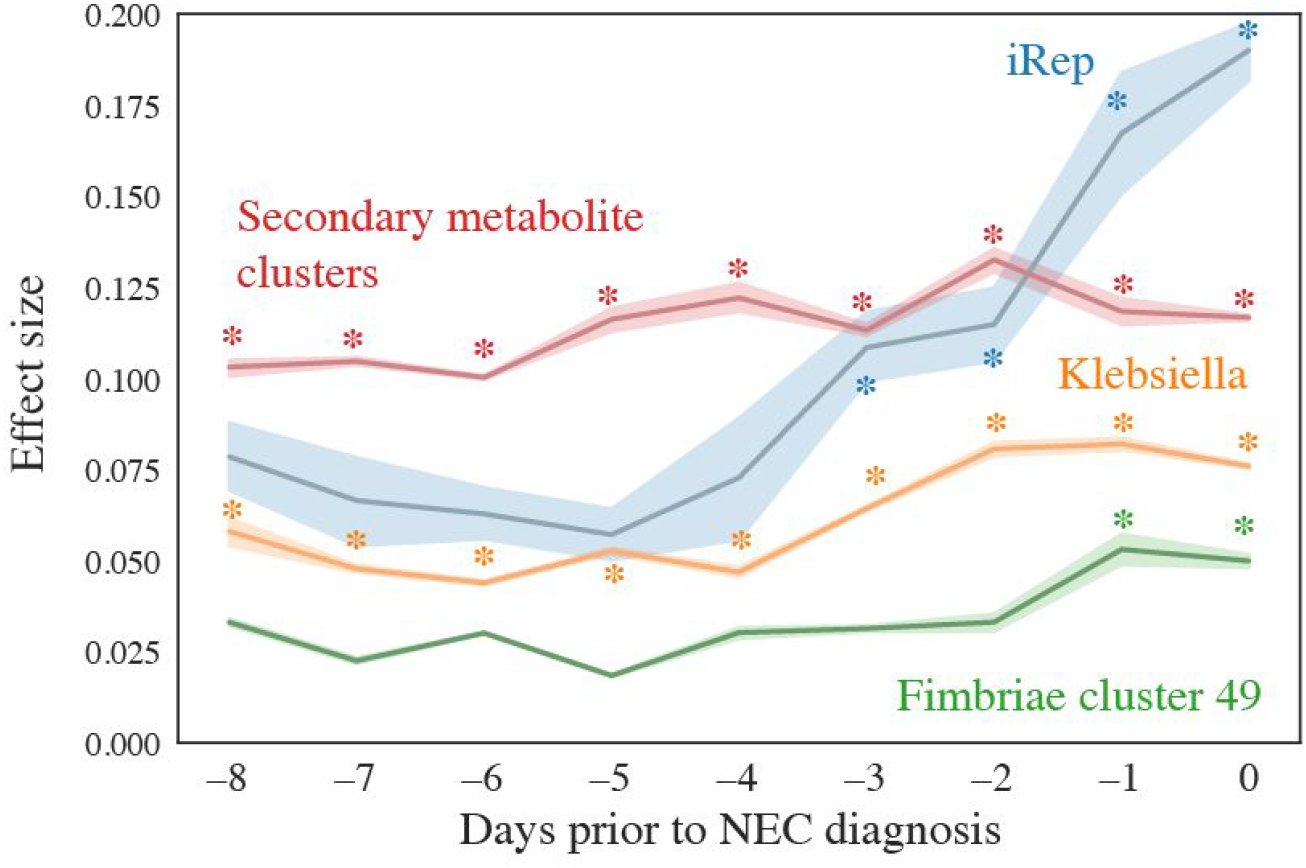
Biomarkers of NEC are most informative closer to NEC diagnosis. The effect size for difference of each feature in pre-NEC vs. control samples is shown based on a Mann Whitney rank sums test over a 2 day sliding window (e.g., −5 compares samples collected from −6 to −4 days relative to NEC diagnosis to control samples). Comparisons with *p* < 0.05 are marked with an asterisk.

## Discussion

Given that we found no single predictor of NEC and identified several factors as important by machine learning, our results support prior indications that NEC is a complex and likely multifactorial disease^2,26^. Of the four aspects of the gut microbiome that differ in pre-NEC compared to control samples (Figure 6), the iRep values of all organisms in each sample had the highest effect size. Given that iRep is a measure of bacterial replication rather than relative abundance, the result highlights that reliance on relative abundance alone could be misleading. This is largely due to the fact that relative abundance metrics are themselves misleading because an organism can increase in relative abundance simply due to the decline in relative abundances of other organisms. The higher bacterial replication rate prior to NEC diagnosis could be sustained by nutrient release from the breakdown of gut tissue. Alternatively, increased bacterial replication may trigger onset of NEC, possibly because high activity of a specific organism leads to imbalance in concentrations of compounds in the gut environment.

Secondary metabolite gene clusters of specific types (bacteriocins, sactipeptides, and butyrolactones) were significantly enriched in pre-NEC compared to control samples (**Supplemental Table S5**). Bacteriocins are small peptides that kill closely related bacteria, and when produced, cell lysis could contribute to onset NEC via release of immunostimulatory compounds such as LPS. Sactipeptides are a class of posttranslationally modified peptides with diverse bioactivities^27^. The sactipeptide with the highest overall importance is related to a subtilosin (antimicrobial agent) with known hemolytic activity. Interestingly, all sactipeptides identified in this study were encoded by Firmicutes, including *Clostridium perfringens* and *Clostridium difficile* (**Supplemental Figure S8; Supplemental Table S5**). Production of sactipeptides by these species could trigger NEC through direct toxicity to human cells or via release of immunostimulatory bacterial compounds following bacterial cell lysis. This phenomenon could explain previous reports that implicate *Clostridium* in development of NEC^28–30^.

Butyrolactones are generally involved in quorum sensing in Actinobacteria^20^, but in this study were mostly found encoded in genomes of Proteobacteria, and over half were identified in *Klebsiella* genomes. Whereas known quorum sensing systems in Proteobacteria are responsible for the production of virulence factors, including fimbriae^31,32^, the functions of butyrolactones in Proteobacteria remain unstudied. Higher proportions of *Klebsiella* were found in infants that went on to develop NEC, and their capacity to produce secondary metabolites and fimbriae could explain this association.

Organisms with genomes encoding for fimbriae cluster 49 were at significantly higher abundances on both the day of and the day before NEC diagnosis. Fimbriae are known stimulants of TLR4 receptors^33^, immune receptors that are overexpressed in premature infants and previously linked to NEC in animal studies^34,35^. Fimbriae are the hallmark pathogenicity factors of uropathogenic *E. coli*^36^, a group of organisms that have been previously implicated as a causative agent of NEC^13^. Uropathogenic *E. coli* were specifically evaluated in this study and not found to be significantly enriched in pre-NEC compared to control samples (**Supplemental Table S3;** Figure 2c). The associations in prior work and the current study may instead reflect a general link between fimbriae and TLR4 receptor stimulation.

An advantage of genome-resolved metagenomics is that it provides whole community information, going far beyond what can be deduced from 16S rRNA gene surveys that are the hallmark of most prior and much current human microbiome research. Here we applied this approach to a sufficiently large dataset to achieve statistical power unprecedented in a genome-resolved metagenomic study, and find that there is there is likely no single bacteriophage, plasmid, eukaryote, virus, or even protein that is responsible for NEC. However, we identify several promising associations through machine learning, many of which have previously been proposed to explain NEC onset but none of which alone can explain all cases. Bacteria of the genus *Klebsiella* emerged from our analyses as organisms of potential importance, with secondary metabolite, LPS and fimbriae production all being possible contributors. The association of these bacteria, as well as bacteria of the *Clostridium* genera, with NEC and their presence in the NICU^19^ supports prior reports proposing that colonization of premature infants by nosocomial microbes may be clinically significant. Overall, we provide insight into how previously proposed but distinct explanations for development of NEC are interconnected, and identify bacterial growth rates as the strongest predictor of disease onset.

## ACKNOWLEDGEMENTS

This research was supported in part by the National Institutes of Health (NIH) under awards RAI092531A and R01-GM109454; the Alfred P. Sloan Foundation under grant APSF-2012-10-05; and National Science Foundation Graduate Research Fellowships to M.O. under Grant No. DGE 1106400. The study was approved by the University of Pittsburgh Institutional Review Board (Protocol PRO10090089). This work used the Vincent J. Coates Genomics Sequencing Laboratory at UC Berkeley, supported by NIH S10 OD018174 Instrumentation Grant.

## Data availability

Reads are available under bioproject under Bioproject PRJNA294605, SRA studies SRP052967, SRP114966, and SRP012558, and SRA accessions SRR5405607 to SRR5406014 (Brooks et al., 2017; Brown et al., 2018; Rahman et al., 2018; RavehSadka et al., 2015, 2016; Sharon et al., 2013)).

## Author Contributions

M.R.O., M.M., and J.F.B. designed the study; M.R.O. performed metagenomic analyses; N.B. and Y.S.S. guided statistical and machine learning analyses, A.C.C. contributed to secondary metabolite analyses; R.B. recruited infants for the study, and B.F. performed all DNA extractions; M.R.O. and J.F.B. wrote the manuscript, and all authors contributed to manuscript revisions.

## METHODS

### Subject recruitment, sample collection, and metagenomic sequencing

This study was reviewed and approved by the University of Pittsburgh Institutional Review Board (IRB PRO12100487 and PRO10090089). This study made use of many different previously analyzed infant datasets. These datasets have previously published descriptions of the study design, patient selection, and sample collection, and are referred to as NIH1^37,38^, NIH2^19^, NIH3^39^, NIH4^40^, NIH5^41^, and Sloan2^19^. Collated sequencing and health information for all infants and samples is provided in the supplemental materials of this manuscript (**Supplemental Tables S1, S2**).

### Metagenomic profiling

#### Read processing and assembly

Reads from all samples were trimmed using Sickle^42^, and reads that mapped to the human genome with Bowtie 2^43^ under default settings were discarded. Reads from all samples were assembled independently using IDBA-UD^44^ under default settings. Co-assemblies were performed for each infant as well, where reads from all samples from that infant were combined and assembled together. Scaffolds <1 kb in length were discarded, and remaining scaffolds were annotated using Prodigal^45^ to predict open reading frames using default metagenomic settings.

#### Recovery of de novo bacterial genomes

DasTool^46^ was used to select the best bacterial bins from the combination of 3 programs for automatic binning-abawaca (https://github.com/CK7/abawaca). concoct^47^, and maxbin2^48^. Cross-mapping was performed between samples for each infant to generate differential abundance signals, and each sample was binned independently. For each infant, dRep v1.4.2^49^ was then used on all bins created from all samples from that infant to generate an infant-specific genome set, using the command “dRep -comp 50 -con 15 --S_algorithm ANImf -sa .99 -nc .25 --checkM_method taxonomy_wf’.

To determine the taxonomy of bins, the amino-acid sequences of all predicted genes were searched against the uniprot database using the command “usearch64 -ublast $aa_file -db uniprot_cp.fasta.udb -maxhits 1 -evalue 0.0001 -threads 6 -blast6out $b6”, and tRep (https://github.com/MrOlm/tRep/tree/master/bin) was used in combination with ETE 3^50^ to convert the list of identified taxIDs into taxonomic levels. Briefly, this assigns a call to each taxonomy level when at least 50% of protein hits reach that taxonomic level.

#### Bacterial growth rates

iRep values^9^ were calculated by first mapping reads from all samples in each infant to the de-replicated genome set from that infant using Bowtie 2. iRep was then run using the command “samtools view $bam | iRep -s - ; iRep_filter.py --long”, and values resulting from genomes with less than 0.9 breadth of coverage were discarded.

To visualize growth rates over time (Figure 3a), all iRep values from all bacteria were averaged together for each DOL relative to NEC and plotted using the seaborn command “sns.pointplot(showfliers=False, ci=68)” (https://seaborn.pydata.org/). Days of life in which less than 5 infants were profiled were manually removed. Boxplots in Figure 3b were also created using seaborn.

#### Bacteriophages, plasmids, and Eukaryotes

For all assemblies, circular contigs were identified using VICA^51^, and bacteriophages were identified using VirSorter^52^ and VirFinder^53^. Bacteriophages were defined as scaffolds that were considered “level 2” or “level 1” by VirSorter, or less than 0.01 p-value by VirFinder. Plasmids were defined as scaffolds which were circular, but not identified as bacteriophage according to the above definition. Bacteriophages and plasmids over 10kb in length were then each de-replicated separately on a per-infant basis using dRep version 2.0.5, with the command “dRep dereplicate -pa .9 --S_algorithm ANImf -nc .5 −l 3000 -N50W 0 -sizeW 1 --noQualityFiltering --clusterAlg singleOverlap”. All plasmid and bacteriophage genomes were then compared to each other using the dRep command “dRep dereplicate -pa .9 --S_algorithm ANImf -nc .5 −l 10000 -N50W 0 -sizeW 1 --noQualityFiltering --clusterAlg single -d”. Eukaryotes were assembled and binned from the gut samples of premature infants as previously reported^41^.

#### Eukaryotic viruses

Eukaryotic viruses were analyzed using the vFam collection^55^, a set of HMMs designed for the identification of eukaryotic viruses within metagenomic sequence data. Hmmsearch^56^ was used to search the HMM set “vFam-A_2014.hmm” against each assembly. All hits with e-values less than 1e-5 were considered significant and retained. Reads were also mapped to a previously curated list of human viruses^57^. This lead to the identification of no viruses when individual samples were used, and a very small number of viruses when combined sets of reads from each infant were used (Torque teno midi virus 2, Torque teno virus 14, and Macaca mulatta polyomavirus 1). This line of work was not followed up on due to lack of signal.

#### Diversity

Shannon diversity and overall bacteria richness were calculated for each sample. Shannon diversity was calculated using the command skbio.diversity.alpha.shannon (http://scikit-bio.org/). Richness was calculated as the number of bacteria with relative abundances over 0.1%.

#### KEGG Modules

KEGG modules were annotated by using HMMER against an in house HMM database built from the Kyoto Encyclopedia of Genes and Genomes (KEGG) orthology groups (KOs)^58^. Briefly, all KEGG database proteins with KOs were compared with all-v-all global similarity search using USEARCH^59^. MCL was then used to sub-cluster KOs (inflation_value = 1.1). Each sub-cluster was aligned using MAFFT^60^, and HMMs were constructed from sub-cluster alignments. HMMs were then scored against all KEGG sequences with KOs and a score threshold was set for each HMM at the score of the highest scoring hit outside of that HMMs sub-cluster. KEGG modules were considered present in a genome if all necessary KOs were present in that genome.

#### Secondary metabolite gene clusters

In order to identify secondary metabolites, antismash-4.0.2 was run on each infant co-assembly^61^. The results were parsed using the custom script parse_antismash.py (https://github.com/MrOlm/Public-Scripts) and resulting key proteins were clustered using diamond^62^ (commands: diamond makedb --in $coassemblies.faa -p 6 -d $coassemblies.faa.db; diamond blastp -q $coassemblies.faa -d $coassemblies.faa.db.dmnd -o $coassemblies.faa.out --id 50). Alignments were filtered to only retain those with >75% amino acid identity and 50% alignment. Hierarchical clustering was then performed using average amino acid identity (AAI) and resolved using a distance threshold of .5 to assign each secondary metabolite gene cluster to a gene cluster family. Next, for each infant the nucleotide sequences of all genes in a representative for each gene cluster family were concatenated together. The reads from each sample from that infant were mapped to this concatenation of genes in order to determine the dynamics of these genes in all samples from that infant. The breadth of each cluster was calculated as the weighted breadth (considering length) for all genes in that cluster.

#### Virulence Factors

Virulence Factors Database (VFDB) was used to search for virulence factors^63^. The database used was from Mar 17, 2017, containing 2597 sequences. Abricate was used to search all predicted protein sequences against the VFDB (https://github.com/tseemann/abricate). A metadata file from VFDB website (http://www.mgc.ac.cn/VFs/) named “FVs.xls” was used to get additional information about the virulence factors. About 15% of virulence factors were not included in this metadata file and were excluded from additional analysis.

#### Botulinum toxin

A blast database of all subtypes of BoNTs toxin was downloaded from from https://bontbase.org/ (as accessed on February 15 2018). Blastp was used to search predicted amino acid sequences of all genes against the database. Hits with an e-value less than 1e-5 were considered valid.

#### Pathogenic E. coli

It was previously reported that pathogenic *E. coli* may be associated with NEC development; specifically the clades 73, 95, 127, 131, 144, 998, abd 69^13^. To identify *E. coli* genomes of these sequencing types in our dataset, all genomes were MLST profiled using PubMLST^64^ and the program “mlst” (https://github.com/tseemann/mlst). The MLST definition requires having 7 genes; in cases where only 6 genes could be identified, if only one sequence type (ST) existed with those 6 gene types, the ST was inferred. Each sample with an *E. coli* genome of the above STs at over 1% relative abundance were considered to have a “pathogenic” *E. coli.*

#### Proteins

Three protein clustering methods were evaluated for use in this study- MMseqs2^65^ (run using default settings), cd-hit (command: “cd-hit -c 0.7 -M 200000 -T 10)^66^, and a previously described hybrid Markov Cluster approach^23^. Algorithms were evaluated based on their ability to reconstruct known protein clusters (specifically a previously described set of 16 universal ribosomal proteins^67^), and the hybrid Markov Cluster approach approach performed best (**Supplemental Table S11**). This method was used to cluster the amino acid sequences of all predicted genes from all assembled scaffolds.

### The average microbiome of NEC and control infants

To calculate the relative abundance of all microbes in each infant, a full “genome inventory” was generated for each infant by resolving the overlap between the recovered bacteria, eukaryote, bacteriophage, and plasmid genomes. Bacteriophage and plasmid genomes were first aligned using nucmer^68^, and in all cases where scaffolds aligned with over 95% ANI on over 50% of the scaffold, the scaffold was removed from the plasmid list. The resulting scaffolds were next aligned to bacterial genomes, and all phage/plasmid scaffolds that aligned to bacterial genomes with the same thresholds were removed. Finally eukaryotic genomes were aligned to the remaining scaffolds, and in cases where similar scaffolds were detected, the scaffold was removed from the eukaryotic genome. Reads from all samples were then mapped to that infant’s genome inventory using Bowtie 2, and the relative abundance of each organism was calculated as the percentage of total samples reads that map to that genome (**Supplemental Table S12**).

In order to compare the microbiome between NEC and control infants, the microbiome of each cohort was averaged across all infants in that cohort (Figure 2a) using the relative abundance values described in the previous paragraph. For each day of life, the average relative abundance of each taxa was first calculated. A 5-day sliding window was next applied, and values from samples in each window were averaged. For example, DOL 10 represents the average abundances from DOL 8 to 12.

### Strain-level differences between NEC and control infants

In order to calculate the relative abundance of each bacterium in each sample, each sample was mapped to the infant-specific bacterial genome set for that infant using Bowtie 2. Relative abundances of all bacteria were calculated as the percentage of total sample reads mapping to each genome (**Supplemental Table S9**). Bacteria assembled from all infants were then compared to each other using dRep, and bacterial genomes with at least 99% ANI were considered to be the same “strain”. A bacterium was considered present in a sample if it had over 0.1% relative abundance, and the fraction of pre-NEC and control samples in which each strain was present was calculated and plotted in Figure 2c.

A similar procedure were performed for the bacteriophage and plasmid genome sets of each infant. Mapping was done to each infant set separately, and genomes were considered to be the same “strain” if they had 99% ANI over at least 50% of their genomes. Organisms were considered present in a sample if they were preset at over 50% genome breadth.

### Principal component analysis

Principal component analysis (PCA) was performed based on the relative abundance of bacteria in each sample (**Supplemental Table S9**) as assessed using weighted UniFrac distance^69^. A phylogenetic tree was created by comparing all assembled bacterial genomes to each other using dRep (command: “dRep --SkipSecondary -ms 100000”), the weighted UniFrac distance between all samples was calculated using scikit-bio (http://scikit-bio.org/), and PCA was performed using scikit-learn^70^.

### Machine learning

#### Algorithm development

2,119 features were calculated for each sample and used as the input to a machine learning classifier to predict pre-NEC and control samples (**Supplemental Table S3**). See above methods for how individual features were calculated.

Three machine learning methods were evaluated for their ability to classify pre-NEC vs. control samples-a random forest classifier (sklearn.ensemble.RandomForestClassifier(n_estimators=460, max_features=10)) balanced using SMOTE (imblearn.combine.SMOTEENN()), a gradient boosting classifier (sklearn.ensemble.GradientBoostingClassifier (learning_rate = 0.1, max_depth = 10, max_features = ‘sqrt’, min_samples_split = 0.7, n_estimators = 200)) balanced using SMOTE, and the same gradient boosting classifier without balancing^70,71^. Hyperparameters were empirically determined using sklearn.model_selection.RandomizedSearchCV, and in general many different combinations of hyperparameters gave similar results. Models were trained and evaluated using cross-validation for 5 iterations each (sklearn.model_selection.StratifiedKFold (n_splits = 10); sklearn.model_selection.cross_val_predict; sklearn.metrics.accuracy_score), and all achieved similar prediction ability (**Supplemental Table S10**).

To determine the accuracy of the gradient boosting classifier, 100 iterations were performed where each iteration consisted of 1) randomly balancing the input to include 21 pre-NEC samples and 21 control samples, 2) classifying each sample in the input using 10 fold cross validation (same methods as above), and 3) calculating the percentage of samples that were correctly classified. The median accuracy value was reported.

#### Feature importance analysis

Feature importances were determined by 100 iterations of training the gradient boosted classifier on the full dataset of pre-NEC and control samples. Importance values were scaled for each iteration such that the overall sum equals 1. The median importance value for each feature is reported (**Supplemental Table S4**).

#### KEGG and secondary metabolite enriched genomes

Each bacterial genome was assigned a metabolic importance value by summing the median feature importances of each KEGG pathway encoded by that genome (see above methods for how KEGG pathways were determined). A distribution of KEGG genomes importances was generated (**Supplemental Figure S4a**), and based on this distribution, genomes with importance values over 15 were considered “organisms of interest”. Each bacterial genome was also assigned an importance value equivalent to the highest importance value of all secondary metabolite clusters encoded by that genome. A distribution was generated (**Supplemental Figure S4b**), and genomes with importances over 0.5 were considered enriched in important secondary metabolite clusters.

### Phylogenetic tree

A phylogenetic tree was made to visualize the distributions of organisms of interest and organisms enriched in important secondary metabolite clusters (Figure 4f). Ribosomal protein S3 was identified in bacterial genomes using pFam PF00189.19 and HMMER with a score cutoff of 50^25,56^. An anchael outgroup was added, and all sequences were aligned using MAFFT^60^ under default parameters. All positions with gaps in over 50% of sequences were trimmed from the alignment, and FastTree was used with default parameters to generate a phylogenetic tree^72^. The tree was visualized and annotated using iTol^73^.

### Protein clustering

#### Protein association with NEC

Each protein cluster was considered present in a sample if a protein from that cluster had been assembled from the sample. Fisher’s exact test was run on each protein cluster to determine if it was enriched in pre-NEC or control samples, and after Benjamini-Hochberg correction no p-values were statistically significant.

#### Protein association with organisms of interest

Each protein cluster was considered present in an organism of interest if a protein from that cluster was encoded in the organism’s genome. The recall and precision of each cluster with organisms of interest was calculated as follows: recall = the number of organisms of interest the cluster in / the total number of organisms of interest; precision = the number of organisms of interest the cluster is in / the total number of genomes the cluster is in. The recall and precision of each protein cluster was plotted (Figure 5b), and clusters with recall and precision over 0.7 were considered enriched in organisms of interest.

The 85 protein clusters enriched in organisms of interest were profiled using using the pFam database^25^ with provided noise cutoffs, and the two most common pFams were PF00419.19 (Fimbrial) and PF00005.26 (ABC transporter) with four proteins each. We next determined if organisms encoding these proteins were enriched in pre-NEC samples. For each pFam with at least three proteins enriched in organisms of interest, we compared the total relative abundance of all bacteria encoding that pFam in pre-NEC vs. control samples, as well as all iRep values of bacteria encoding that pFam in pre-NEC vs. control samples using the Wilcoxon rank-sum test with Benjamini-Hochberg p-value correction (**Supplemental Table S8**).

### Fimbriae

Chaperone-usher fimbriae were identified in our dataset using pFam PF00577.19 (usher protein) and clustered using usearch^59^ with the command “usearch -cluster_fast -id 0.9”. The taxonomic profile of each fimbriae cluster was determined based on the taxonomy of organisms encoded by that cluster, and relative abundance and iRep associations with pre-NEC vs. control samples were calculated using the Wilcoxon rank-sum test applied to all bacterial genomes encoding each cluster. A similar procedure was performed using genomes which were not classified as organisms of interest but did encode fimbriae cluster 49, comparing between pre-NEC and control samples and between all samples from NEC infants and all samples from control infants (**Supplemental Figure S6**).

A phylogenetic tree was made in order to establish the type of usher proteins identified in our study. Three reference sequences from each previously established type^24^ were aligned with three representatives of each of our clusters using MAFFT. All columns with gaps in over 50% of sequences were trimmed from the alignment, IQtree was used with default parameters to generate a phylogenetic tree^74^, and tree annotation was performed using iTol^73^.

### Effect size calculations

To determine when signals first become apparent relative to NEC diagnosis, control samples were compared to samples collected over different sliding 3-day windows (Figure 6). To compare the signal at 5 days prior to NEC diagnosis, for example, a rarefied set of samples was chosen from 4-6 days prior to diagnosis where one sample from each infant that has a sample in that window was randomly chosen. This procedure was repeated 10 times, and the average effect size and 95% confidence intervals were plotted. The effect size was calculated based on the Wilcoxon rank-sum test statistic (as calculated by SciPy (scipy.stats.ranksums)^75^) using the formula: effect size = (test statistic / square root ((observations in population 1) + (observations in population 2))). For *iRep* all iRep values were compared between the two sets, for *secondary metabolite gene clusters* the total relative abundance of genomes encoding secondary metabolite gene clusters classified as producing sactipeptides, bacteriocins, or butyrolactones was compared, for *Klebsiella* the total relative abundances of all genomes classified as the genus *Klebsiella* were compared, and for *Fimbriae cluster 49* the total relative abundances of all genomes encoding fimbriae cluster 49 were compared.

## REFERENCES

1. Obladen, M. Necrotizing enterocolitis--150 years of fruitless search for the cause. Neonatology 96, 203–210 (2009).

2. Neu, J. & Walker, W. A. Necrotizing enterocolitis. N. Engl. J. Med. 364, 255–264 (2011).

3. Claud, E. C. et al. Developmentally regulated IκB expression in intestinal epithelium and susceptibility to flagellin-induced inflammation. Proc. Natl. Acad. Sci. U. S. A. 101, 7404–7408 (2004).

4. Denning, N.-L. & Prince, J. M. Neonatal intestinal dysbiosis in necrotizing enterocolitis. Mol. Med. 24, 4 (2018).

5. Afrazi, A. et al. New insights into the pathogenesis and treatment of necrotizing enterocolitis: Toll-like receptors and beyond. Pediatr. Res. 69, 183–188 (2011).

6. Lawrence, G., Bates, J. & Gaul, A. PATHOGENESIS OF NEONATAL NECROTISING ENTEROCOLITIS. Lancet 319, 137–139 (1982).

7. Hosny, M., Cassir, N. & La Scola, B. Updating on gut microbiota and its relationship with the occurrence of necrotizing enterocolitis. Human Microbiome Journal 4, 14–19 (2017).

8. Pammi, M. et al. Intestinal dysbiosis in preterm infants preceding necrotizing enterocolitis: a systematic review and meta-analysis. Microbiome 5, (2017).

9. Brown, C. T., Olm, M. R., Thomas, B. C. & Banfield, J. F. Measurement of bacterial replication rates in microbial communities. Nat. Biotechnol. 34, 1256–1263 (2016).

10. Korem, T. et al. Growth dynamics of gut microbiota in health and disease inferred from single metagenomic samples. Science (2015). doi:10.1126/science.aac4812

11. Brown, C. T. et al. Unusual biology across a group comprising more than 15% of domain Bacteria. Nature 523, 208–211 (2015).

12. Ronda, C., Chen, S. P., Cabral, V., Yaung, S. J. & Wang, H. H. Metagenomic engineering of the mammalian gut microbiome in situ. Nat. Methods (2019). doi: 10.1038/s41592-018-0301-y

13. Ward, D. V. et al. Metagenomic Sequencing with Strain-Level Resolution Implicates Uropathogenic E. coli in Necrotizing Enterocolitis and Mortality in Preterm Infants. Cell Rep. (2016). doi:10.1016/j.celrep.2016.03.015

14. Zhang, S. et al. Identification of a Botulinum Neurotoxin-like Toxin in a Commensal Strain of Enterococcus faecium. Cell Host Microbe 23, 169–176.e6 (2018).

15. Penders, J. et al. Factors Influencing the Composition of the Intestinal Microbiota in Early Infancy. Pediatrics 118, 511–521 (2006).

16. Bokulich, N. A. et al. Antibiotics, birth mode, and diet shape microbiome maturation during early life. Sci. Transl. Med. 8, 343ra82–343ra82 (2016).

17. Khan, H. A., Baig, F. K. & Mehboob, R. Nosocomial infections: Epidemiology, prevention, control and surveillance. Asian Pac. J. Trop. Biomed. 7, 478–482 (2017).

18. Morrow, A. L. et al. Early microbial and metabolomic signatures predict later onset of necrotizing enterocolitis in preterm infants. Microbiome 1, 1 (2013).

19. Brooks, B. et al. Strain-resolved analysis of hospital rooms and infants reveals overlap between the human and room microbiome. Nat. Commun. 8, 1814 (2017).

20. Du, Y.-L., Shen, X.-L., Yu, P., Bai, L.-Q. & Li, Y.-Q. Gamma-butyrolactone regulatory system of Streptomyces chattanoogensis links nutrient utilization, metabolism, and development. Appl. Environ. Microbiol. 77, 8415–8426 (2011).

21. Huang, T. et al. Isolation of a variant of subtilosin A with hemolytic activity. J. Bacteriol. 191, 5690–5696 (2009).

22. Miyamoto, K. et al. Complete sequencing and diversity analysis of the enterotoxin-encoding plasmids in Clostridium perfringens type A non-food-borne human gastrointestinal disease isolates. J. Bacteriol. 188, 1585–1598 (2006).

23. Meheust, R., Burstein, D., Castelle, C. J. & Banfield, J. F. Biological capacities clearly define a major subdivision in Domain Bacteria. bioRxiv 335083 (2018). doi: 10.1101/335083

24. Nuccio, S.-P. & Bäumler, A. J. Evolution of the chaperone/usher assembly pathway: fimbrial classification goes Greek. Microbiol. Mol. Biol. Rev. 71, 551–575 (2007).

25. El-Gebali, S. et al. The Pfam protein families database in 2019. Nucleic Acids Res. (2018). doi: 10.1093/nar/gky995

26. Ballance, W. A., Dahms, B. B., Shenker, N. & Kliegman, R. M. Pathology of neonatal necrotizing enterocolitis: a ten-year experience. J. Pediatr. 117, S6–13 (1990).

27. Arnison, P. G. et al. Ribosomally synthesized and post-translationally modified peptide natural products: overview and recommendations for a universal nomenclature. Nat. Prod. Rep. 30, 108–160 (2013).

28. Morowitz, M. J., Poroyko, V., Caplan, M., Alverdy, J. & Liu, D. C. Redefining the role of intestinal microbes in the pathogenesis of necrotizing enterocolitis. Pediatrics 125, 777–785 (2010).

29. de la Cochetiere, M.-F. et al. Early intestinal bacterial colonization and necrotizing enterocolitis in premature infants: the putative role of Clostridium. Pediatr. Res. 56, 366–370 (2004).

30. Dittmar, E. et al. Necrotizing enterocolitis of the neonate with Clostridium perfringens: diagnosis, clinical course, and role of alpha toxin. Eur. J. Pediatr. 167, 891–895 (2008).

31. Sturbelle, R. T. et al. The role of quorum sensing in Escherichia coli (ETEC) virulence factors. Vet. Microbiol. 180, 245–252 (2015).

32. Rutherford, S. T. & Bassler, B. L. Bacterial quorum sensing: its role in virulence and possibilities for its control. Cold Spring Harb. Perspect. Med. 2, (2012).

33. Fischer, H., Yamamoto, M., Akira, S., Beutler, B. & Svanborg, C. Mechanism of pathogen-specific TLR4 activation in the mucosa: fimbriae, recognition receptors and adaptor protein selection. Eur. J. Immunol. 36, 267–277 (2006).

34. Leaphart, C. L. et al. A critical role for TLR4 in the pathogenesis of necrotizing enterocolitis by modulating intestinal injury and repair. J. Immunol. 179, 4808–4820 (2007).

35. Jilling, T. et al. The roles of bacteria and TLR4 in rat and murine models of necrotizing enterocolitis. J. Immunol. 177, 3273–3282 (2006).

36. Wiles, T. J., Kulesus, R. R. & Mulvey, M. A. Origins and virulence mechanisms of uropathogenic Escherichia coli. Exp. Mol. Pathol. 85, 11–19 (2008).

37. Brown, C. T. et al. Hospitalized Premature Infants Are Colonized by Related Bacterial Strains with Distinct Proteomic Profiles. MBio 9, (2018).

38. Raveh-Sadka, T. et al. Evidence for persistent and shared bacterial strains against a background of largely unique gut colonization in hospitalized premature infants. ISME J. (2016). doi: 10.1038/ismej.2016.83

39. Raveh-Sadka, T. et al. Gut bacteria are rarely shared by co-hospitalized premature infants, regardless of necrotizing enterocolitis development. Elife 4, e05477 (2015).

40. Rahman, S. F., Olm, M. R., Morowitz, M. J. & Banfield, J. F. Machine learning leveraging genomes from metagenomes identifies influential antibiotic resistance genes in the infant gut microbiome. mSystems 3, e00123–17 (2018).

41. Olm, M. R. et al. Strain-level overlap between infant and hospital fungal microbiomes revealed through de novo assembly of eukaryotic genomes from metagenomes. bioRxiv 324566 (2018). doi: 10.1101/324566

42. Joshi, N. A. & Fass, J. N. Sickle: a sliding-window, adaptive, quality-based trimming tool for FastQ files. Available from: github.com/najoshi/sickle (2011).

43. Langmead, B. & Salzberg, S. L. Fast gapped-read alignment with Bowtie 2. Nat. Methods 9, 357–359 (2012).

44. Peng, Y., Leung, H. C. M., Yiu, S. M. & Chin, F. Y. L. IDBA-UD: a de novo assembler for single-cell and metagenomic sequencing data with highly uneven depth. Bioinformatics 28, 1420–1428 (2012).

45. Hyatt, D. et al. Prodigal: prokaryotic gene recognition and translation initiation site identification. BMC Bioinformatics 11, 119 (2010).

46. Sieber, C. M. K. et al. Recovery of genomes from metagenomes via a dereplication, aggregation and scoring strategy. Nature Microbiology 3, 836–843 (2018).

47. Alneberg, J. et al. Binning metagenomic contigs by coverage and composition. Nat. Methods 11, 1144–1146 (2014).

48. Wu, Y.-W., Simmons, B. A. & Singer, S. W. MaxBin 2.0: an automated binning algorithm to recover genomes from multiple metagenomic datasets. Bioinformatics 32, 605–607 (2016).

49. Olm, M. R., Brown, C. T., Brooks, B. & Banfield, J. F. dRep: a tool for fast and accurate genomic comparisons that enables improved genome recovery from metagenomes through de-replication. ISME J. (2017). doi: 10.1038/ismej.2017.126

50. Huerta-Cepas, J., Serra, F. & Bork, P. ETE 3: Reconstruction, Analysis, and Visualization of Phylogenomic Data. Mol. Biol. Evol. 33, 1635–1638 (2016).

51. Crits-Christoph, A., Gelsinger, D. R. & Ma, B. Functional interactions of archaea, bacteria and viruses in a hypersaline endolithic community. Environmentalist (2016).

52. Roux, S., Enault, F., Hurwitz, B. L. & Sullivan, M. B. VirSorter: mining viral signal from microbial genomic data. PeerJ 3, e985 (2015).

53. Ren, J., Ahlgren, N. A., Lu, Y. Y., Fuhrman, J. A. & Sun, F. VirFinder: a novel k-mer based tool for identifying viral sequences from assembled metagenomic data. Microbiome 5, (2017).

54. Zaharia, M. et al. Faster and more accurate sequence alignment with SNAP. *arXivpreprint arXiv:*1111. 5572 (2011).

55. Skewes-Cox, P., Sharpton, T. J., Pollard, K. S. & DeRisi, J. L. Profile hidden Markov models for the detection of viruses within metagenomic sequence data. PLoS One 9, e105067 (2014).

56. Finn, R. D., Clements, J. & Eddy, S. R. HMMER web server: interactive sequence similarity searching. Nucleic Acids Res. 39, (2011).

57. Rampelli, S. et al. ViromeScan: a new tool for metagenomic viral community profiling. BMC Genomics 17, 165 (2016).

58. Kanehisa, M. et al. Data, information, knowledge and principle: back to metabolism in KEGG. Nucleic Acids Res. 42, D199–D205 (2014).

59. Edgar, R. C. Search and clustering orders of magnitude faster than BLAST. Bioinformatics 26, 2460–2461 (2010).

60. Katoh, K. & Standley, D. M. MAFFT multiple sequence alignment software version 7: improvements in performance and usability. Mol. Biol. Evol 30, 772–780 (2013).

61. Weber, T. et al. antiSMASH 3.0–a comprehensive resource for the genome mining of biosynthetic gene clusters. Nucleic Acids Res. 43, W237–W243 (2015).

62. Buchfink, B., Xie, C. & Huson, D. H. Fast and sensitive protein alignment using DIAMOND. Nat. Methods 12, 59–60 (2015).

63. Chen, L. et al. VFDB: a reference database for bacterial virulence factors. Nucleic Acids Res. 33, D325–8 (2005).

64. Jolley, K. A. & Maiden, M. C. J. BIGSdb: Scalable analysis of bacterial genome variation at the population level. BMC Bioinformatics 11, 595 (2010).

65. Steinegger, M. & Söding, J. MMseqs2 enables sensitive protein sequence searching for the analysis of massive data sets. Nat. Biotechnol. 35, 1026–1028 (2017).

66. Huang, Y., Niu, B., Gao, Y., Fu, L. & Li, W. CD-HIT Suite: a web server for clustering and comparing biological sequences. Bioinformatics 26, 680–682 (2010).

67. Hug, L. A. et al. A new view of the tree of life. Nat Microbiol 1, 16048 (2016).

68. Delcher, A. L., Salzberg, S. L. & Phillippy, A. M. Using MUMmer to identify similar regions in large sequence sets. Curr. Protoc. Bioinformatics Chapter 10, Unit 10.3 (2003).

69. Lozupone, C. & Knight, R. UniFrac: a New Phylogenetic Method for Comparing Microbial Communities. Appl. Environ. Microbiol. 71, 8228–8235 (2005).

70. Pedregosa, F. et al. Scikit-learn: Machine learning in Python. J. Mach. Learn. Res. 12, 2825–2830 (2011).

71. Chawla, N. V., Bowyer, K. W., Hall, L. O. & Kegelmeyer, W. P. SMOTE: Synthetic Minority Over-sampling Technique. 1 16, 321–357 (2002).

72. Price, M. N., Dehal, P. S. & Arkin, A. P. FastTree: computing large minimum evolution trees with profiles instead of a distance matrix. Mol. Biol. Evol. 26, (2009).

73. Letunic, I. & Bork, P. Interactive Tree Of Life (iTOL): an online tool for phylogenetic tree display and annotation. Bioinformatics 23, 127–128 (2007).

74. Nguyen, L.-T., Schmidt, H. A., von Haeseler, A. & Minh, B. Q. IQ-TREE: a fast and effective stochastic algorithm for estimating maximum-likelihood phylogenies. Mol. Biol. Evol 32, 268–274 (2015).

75. Jones, E., Oliphant, T. & Peterson, P. SciPy: Open source scientific tools for Python. URL http://scipy.org (2001).

