## Supplemental Captions and Figures for "Necrotizing enterocolitis is preceded by increased gut bacterial replication, *Klebsiella*, and fimbriae-encoding bacteria that may stimulate TLR4 receptors"

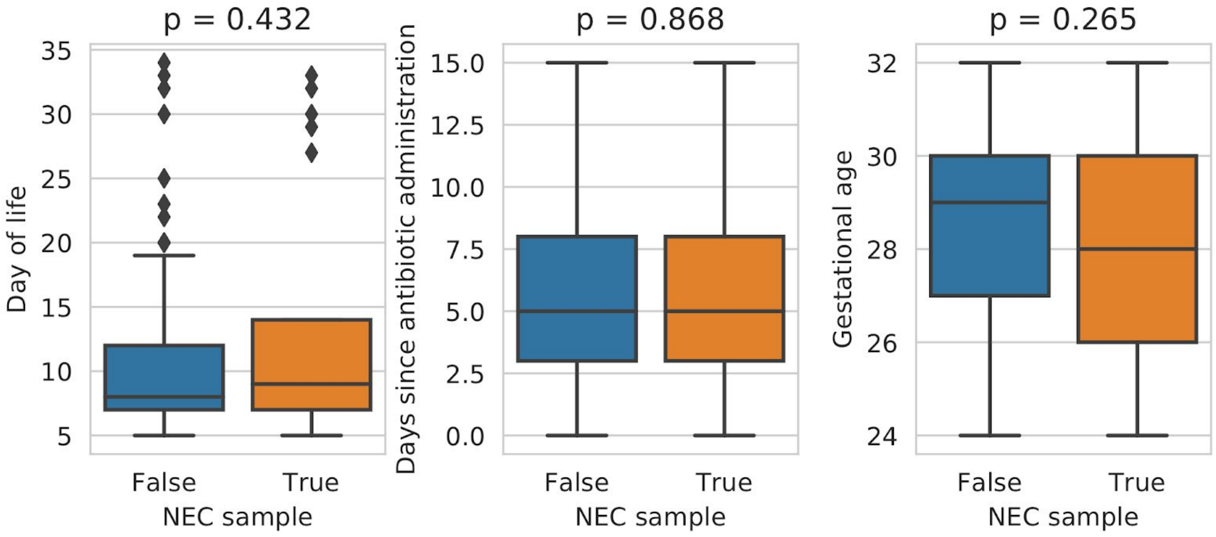


**Supplemental Figure S1**: Distribution of clinical metadata in metagenomic samples used for statistical tests. There are 21 pre-NEC samples (all within 2 days of NEC diagnosis) and 126 control samples. Distributions were compared using the Wilcoxon rank sums test, with the overall p-value reported.

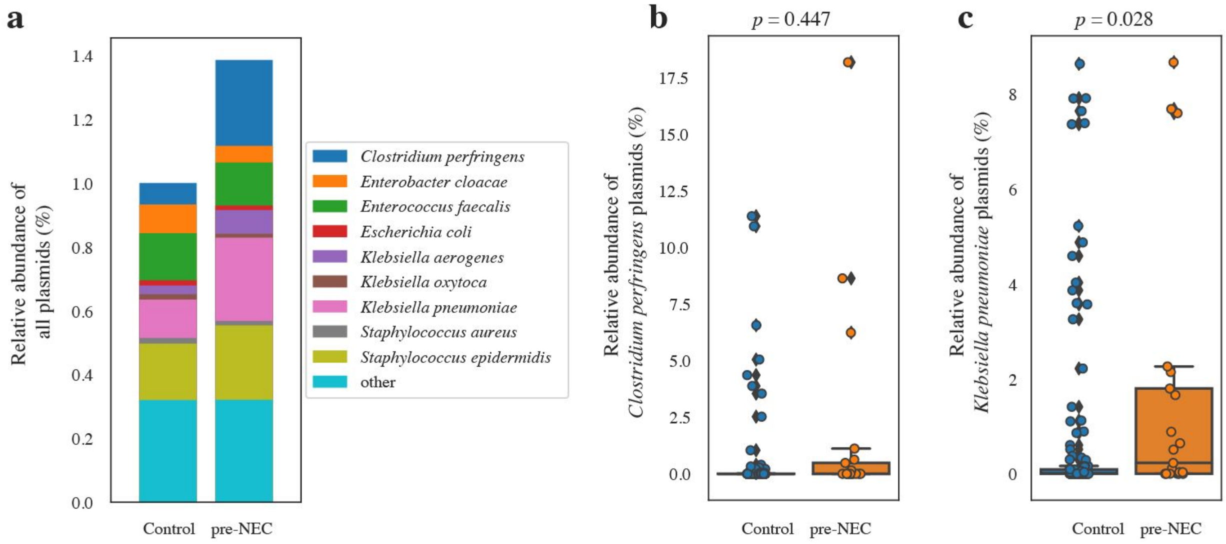


**Supplemental Figure S3:** Total plasmid content in pre-NEC vs. control samples. (**a**) Taxonomic distribution of plasmids in control and pre-NEC samples. Height of bars represents the average relative abundance of plasmids of each species-level taxa. (**b, c**) Difference in the total abundance of *C. perfringens* plasmids (**b**) and *Klebsiella pneumoniae* (**c**) plasmids in control and pre-NEC samples. *P*-values listed above each plot are from Wilcoxon rank-sums test.

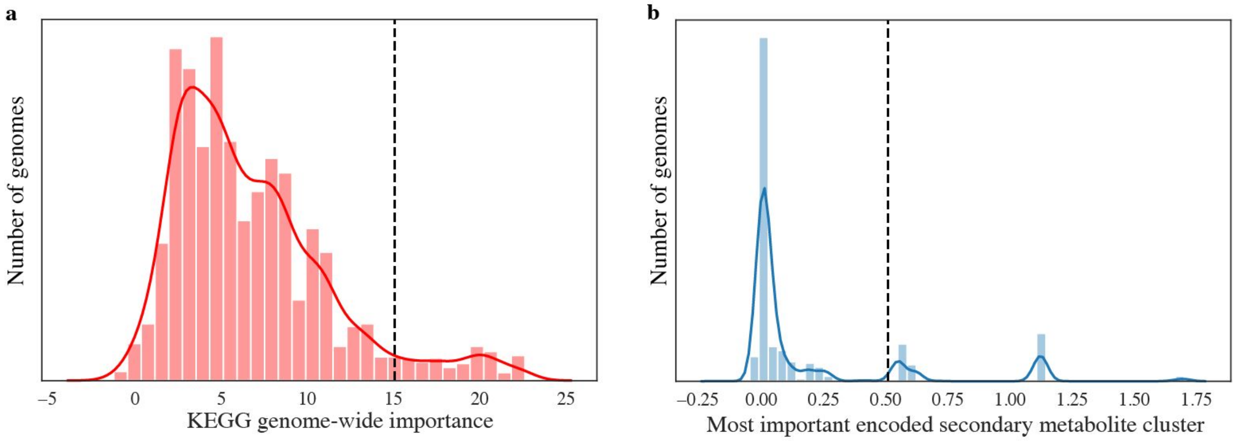


**Supplemental Figure S4**: Genome-wide associations with important KEGG modules and important secondary metabolite clusters. (**a**) The distribution of total genome KEGG module importances. Those with overall importances above 15 were considered "metabolically important. Total genome KEGG module importance was calculated by taking the sum of the importances of all KEGG modules encoded by each genome. (**b**) The distribution of the most important secondary metabolite cluster of each genome. Genomes without a secondary metabolite cluster are not included. Genomes encoding a secondary metabolite cluster with an importance over 0.5 were to encode important secondary metabolite clusters.

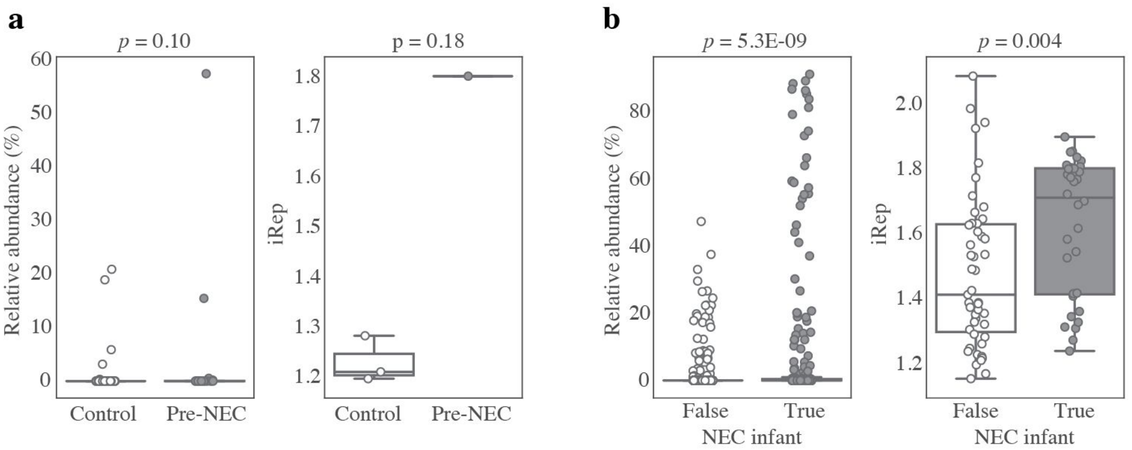


**Supplemental Figure S6**: NEC association with genomes which are not organisms of interest but encode fimbriae cluster 49. (**a**) Comparing the abundance and iRep of these organisms in pre-NEC and control infants does not achieve statistical significance, seemingly because there are not enough data-points to compare. (**b**) Comparing the abundance and iRep of these organisms in all samples from NEC infants vs. all samples from control infants does achieve statistical significance. P-values from Wilcoxon rank-sums test.

(LARGE PDF)

**Supplemental Figure S7:** Visualization of metadata as it relates to principal component analysis. The first sheet shows pre-NEC (red) and control (black) samples across the top 5 principal components. Subsequent sheets show the the first two principal components of both matched and all samples, colored by 8 different pieces of health metadata.

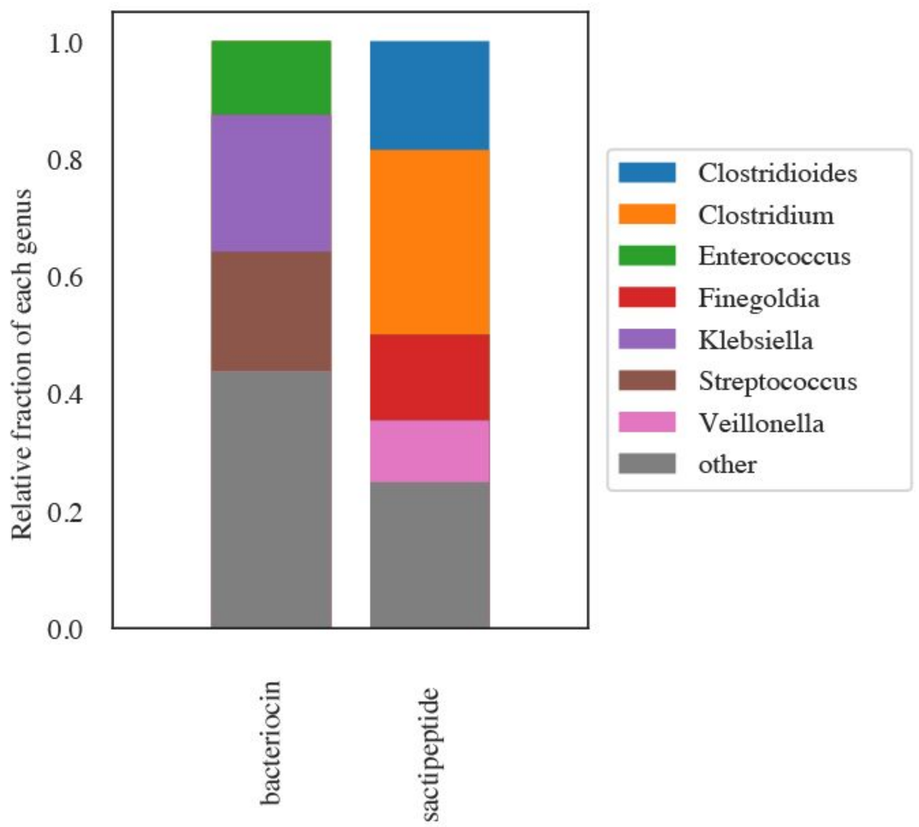


**Supplemental Figure S8**: Genus-level taxonomic makeup of genomes encoding two types of secondary metabolite clusters enriched in NEC infants. Taxa at less than 6% relative abundance are put into the "other" category.

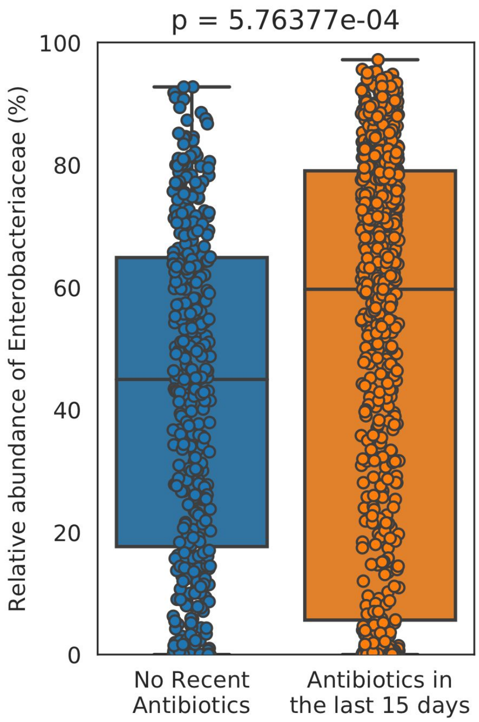


**Supplemental Figure S9**: Relative abundance of Enterobacteriaceae in samples within 15 days of antibiotic administration vs. not. *P*-value from Wilcoxon rank-sums test.

**Supplemental Table S1**: Metagenomic sequencing information for all samples

**Supplemental Table S2**: Patient metadata

**Supplemental Table S3**: Full list of metadata provided to the machine learning classifier. Feature names are coded using the format "category $ type of data $ value"

**Supplemental Table S4**: Importance of all features resulting from the machine learning classifier

**Supplemental Table S5**: Information about de-replicated secondary metabolite clusters

**Supplemental Table S6**: Information on *de novo* assembled genomes

**Supplemental Table S7**: Genome-wide importance of genomes based on secondary metabolites and KEGG modules

**Supplemental Table S8**: Proteins enriched in genomes of interest

**Supplemental Table S9**: Relative abundance and iRep values of genomes in each sample

**Supplemental Table S10**: Accuracy of machine learning algorithms

**Supplemental Table S11**: Accuracy of protein clustering algorithms

**Supplemental Table S12**: Averaged abundance of taxa in NEC and control infants

**Supplemental Table S13**: Identified fimbrial genes
