## Supplemental Figure S7 for "Necrotizing enterocolitis is preceded by increased gut bacterial replication, *Klebsiella*, and fimbriae-encoding bacteria that may stimulate TLR4 receptors"

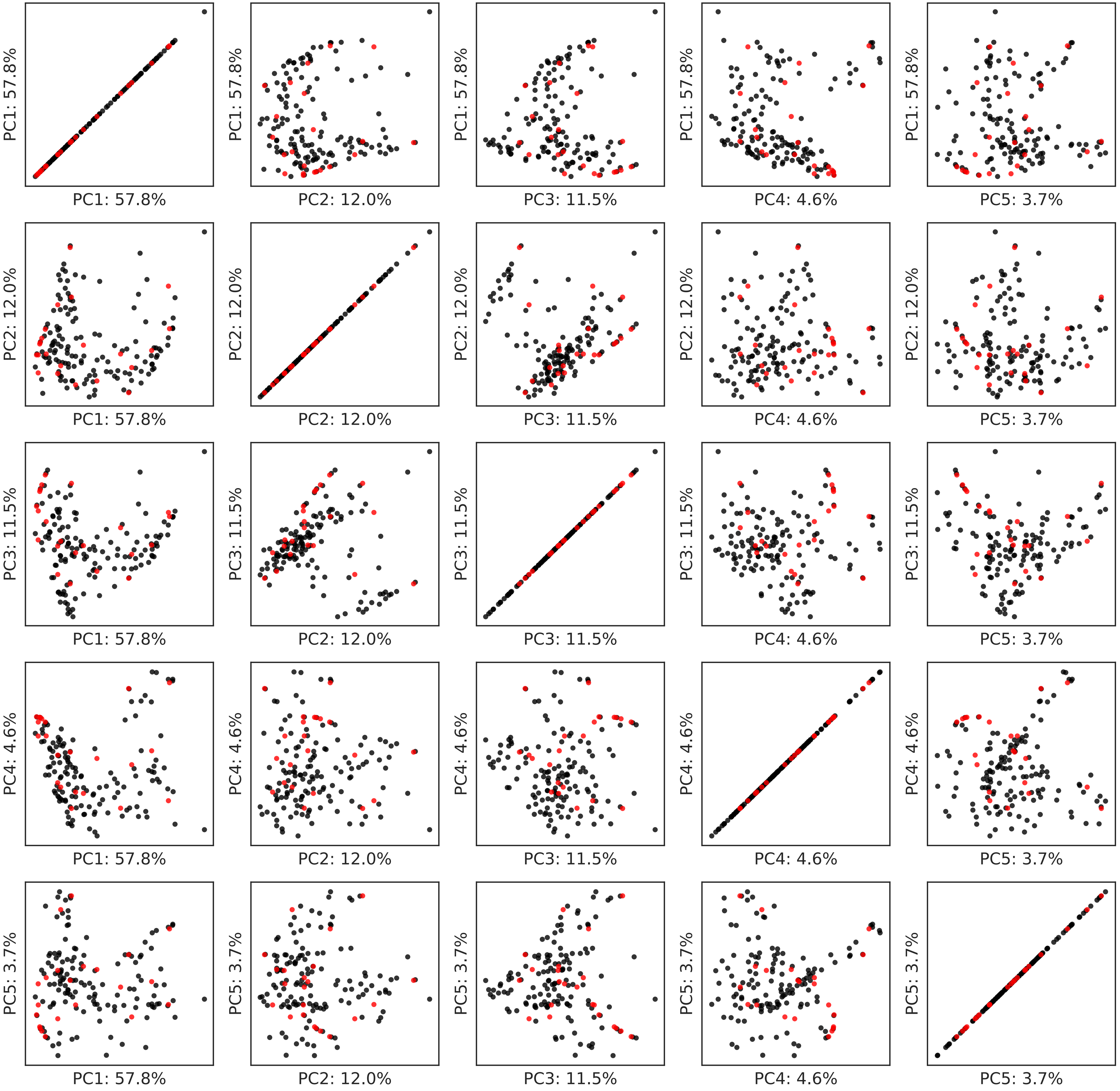

PC1: 57.8%
PC2: 12.0%
PC3: 11.5%
PC4: 4.6%
PC5: 3.7%

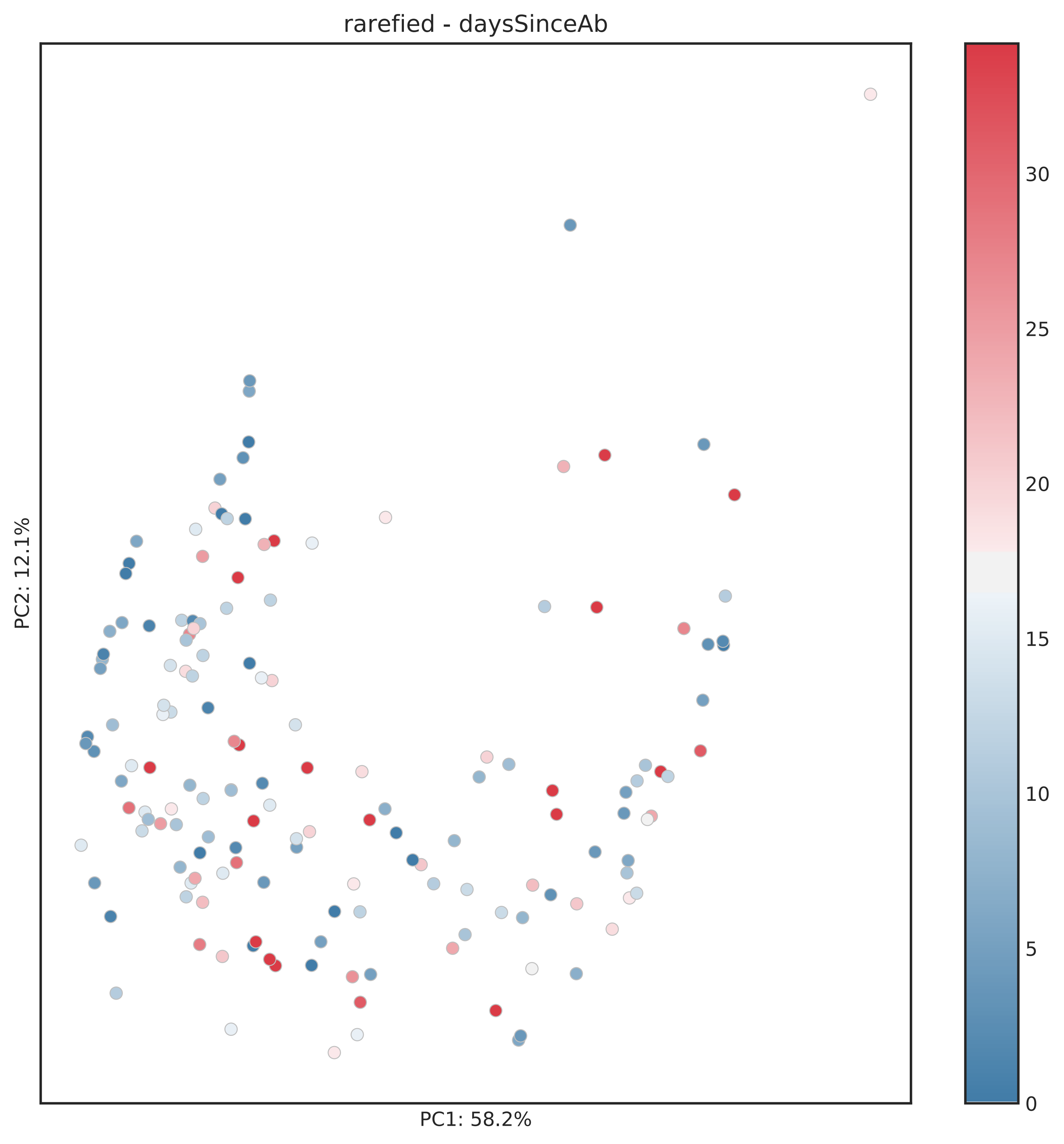
rarefied - daysSinceAb
PC1: 58.2%
PC2: 12.1%
0
5
10
15
20
25
30

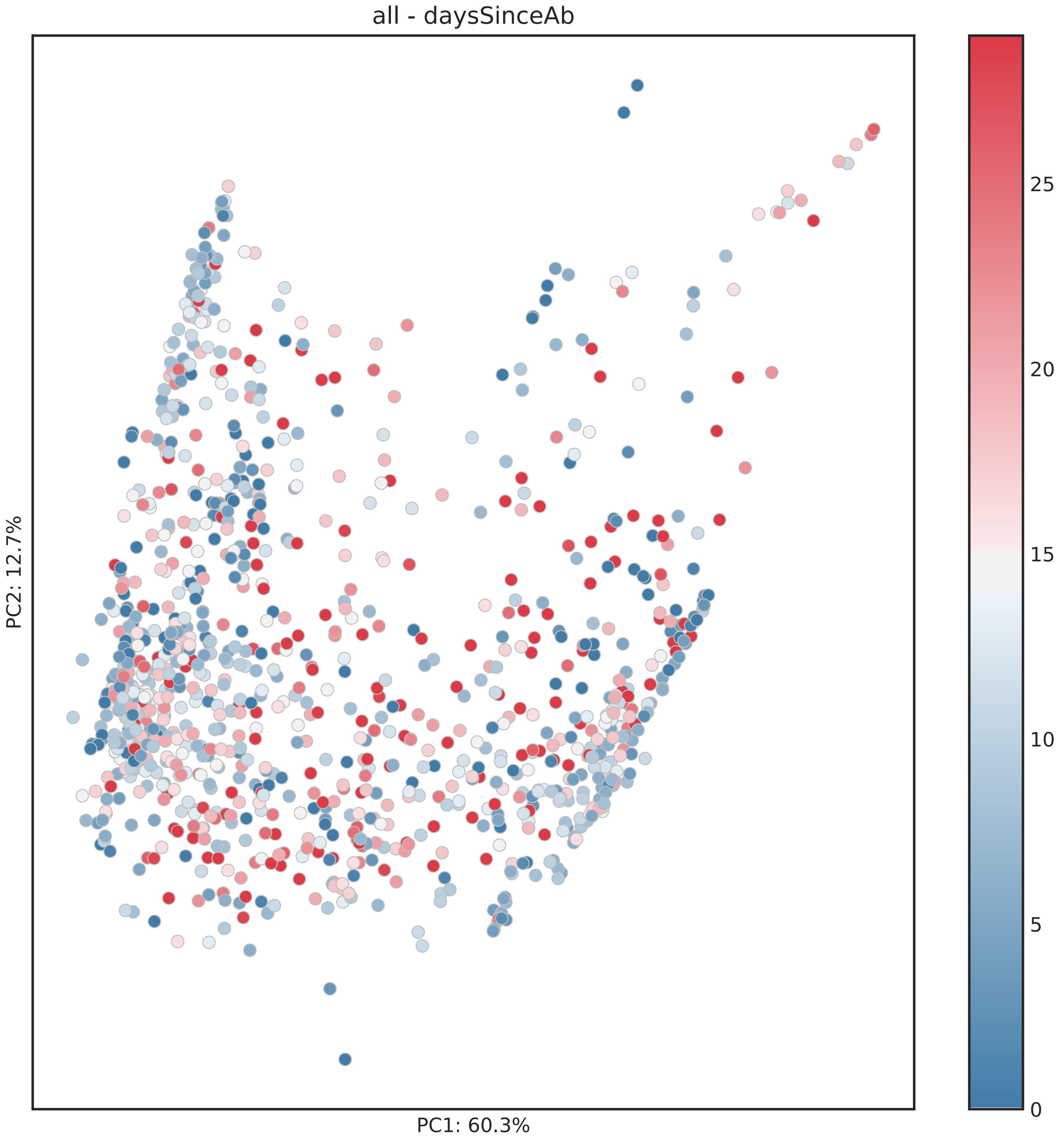

all - daysSinceAb
PC2: 12.7%
PC1: 60.3%
25
20
15
10
5
0

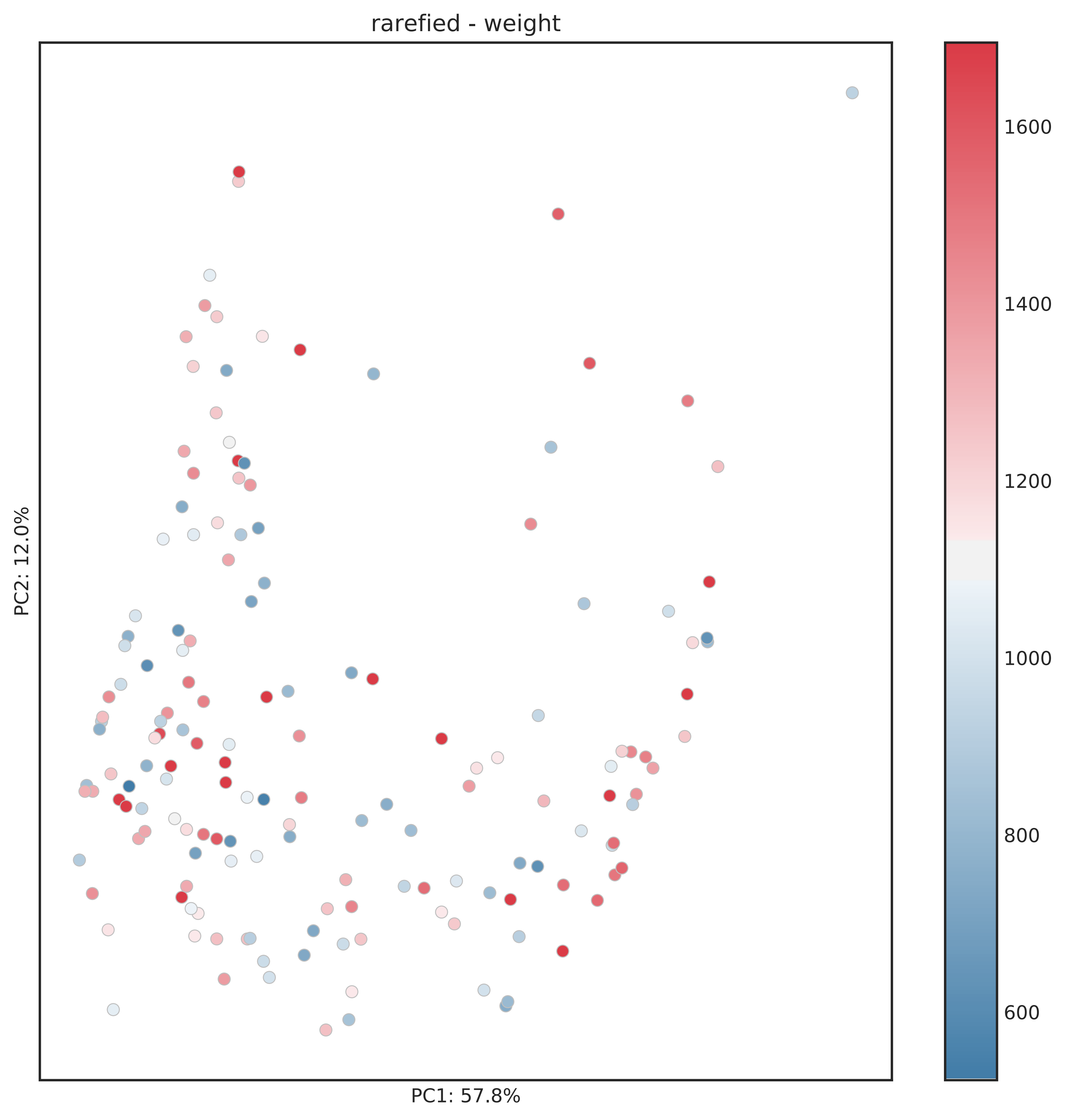
rarefied - weight
PC1: 57.8%
PC2: 12.0%
1600
1400
1200
1000
800
600

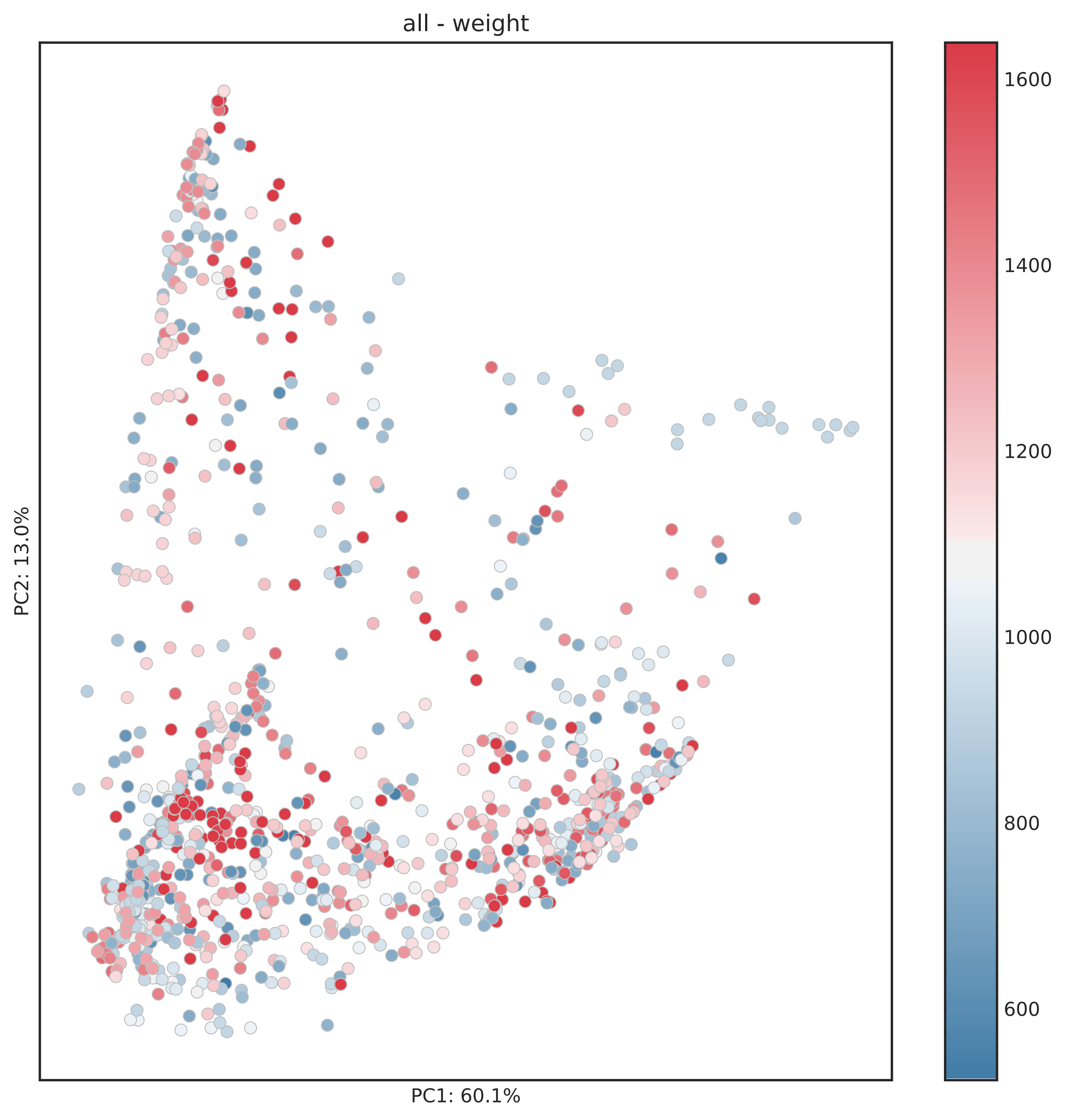
all - weight
PC1: 60.1%
PC2: 13.0%
1600
1400
1200
1000
800
600

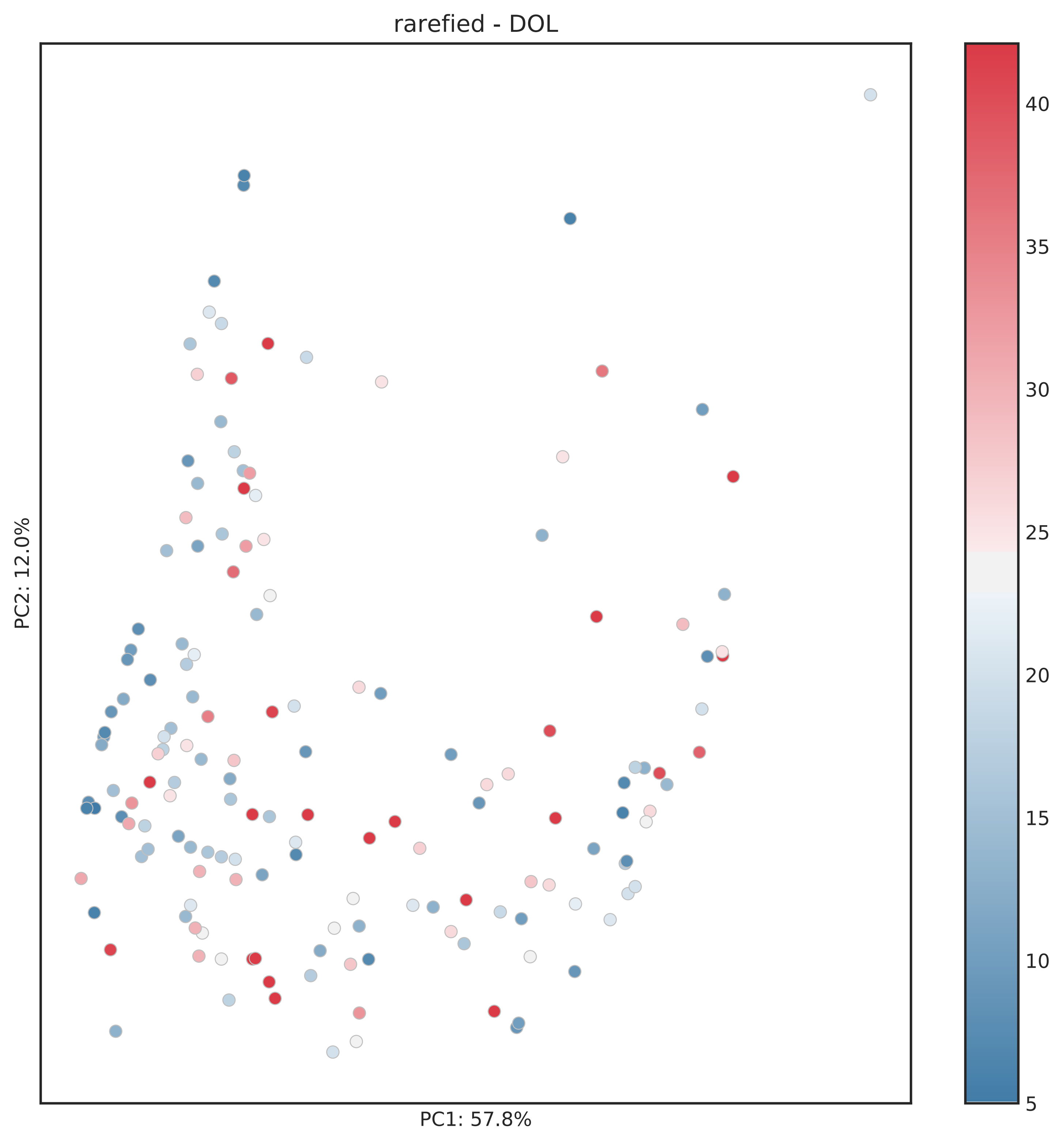
rarefied - DOL
PC1: 57.8%
PC2: 12.0%
5
10
15
20
25
30
35
40

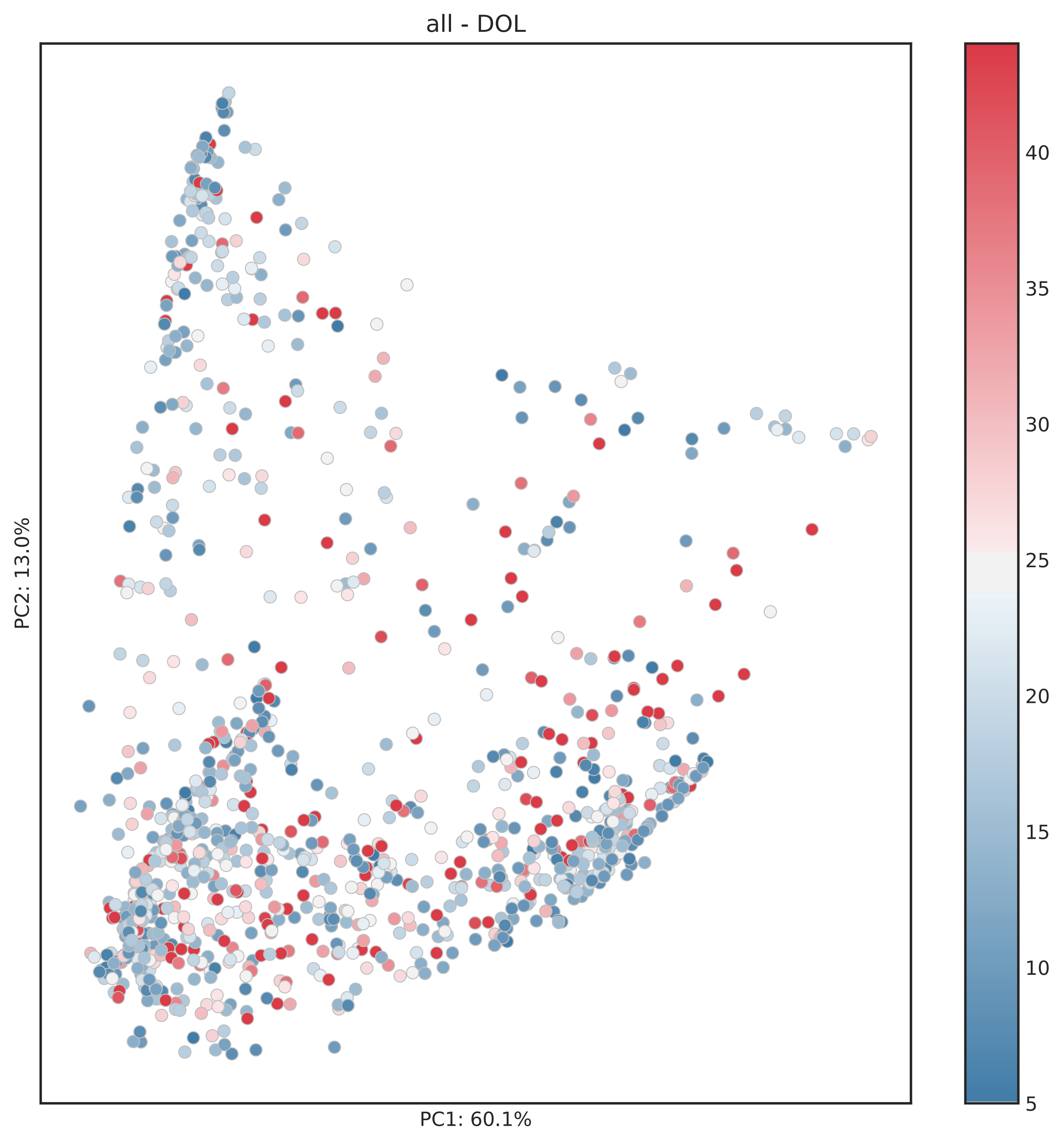
all - DOL
PC1: 60.1%
PC2: 13.0%
5
10
15
20
25
30
35
40

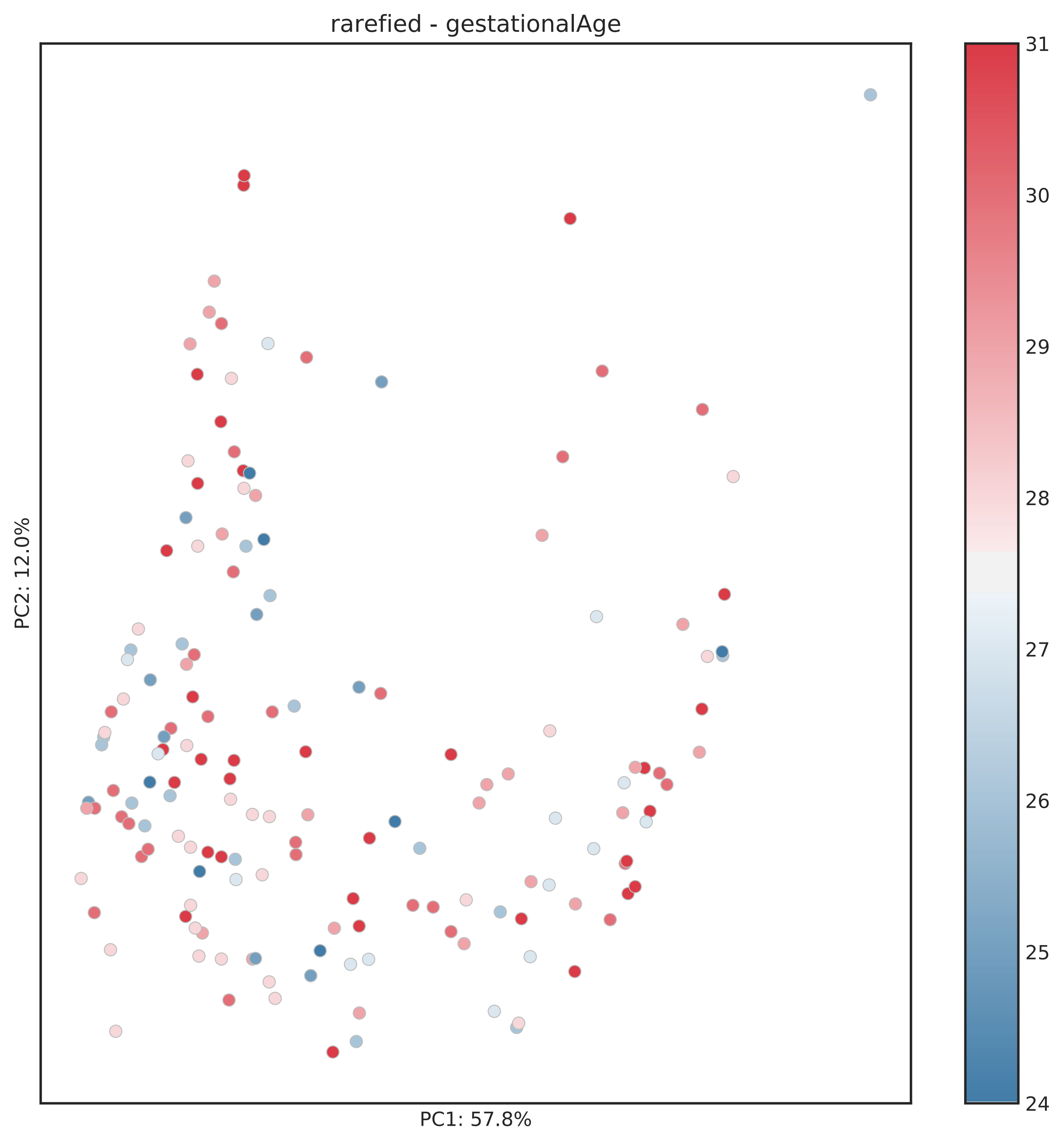
rarefied - gestationalAge
PC1: 57.8%
PC2: 12.0%
31
30
29
28
27
26
25
24

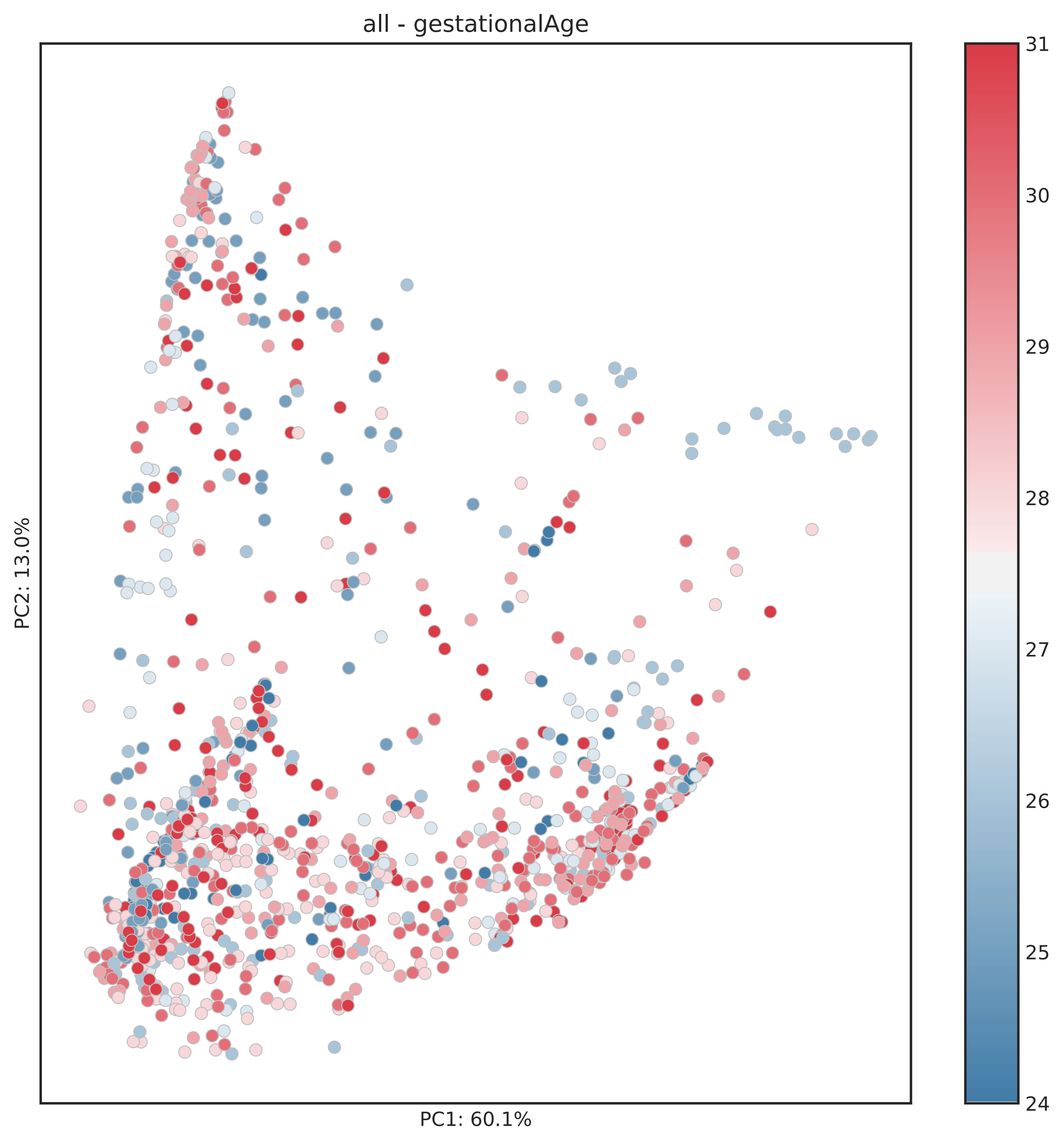
all - gestationalAge
PC2: 13.0%
PC1: 60.1%
31
30
29
28
27
26
25
24

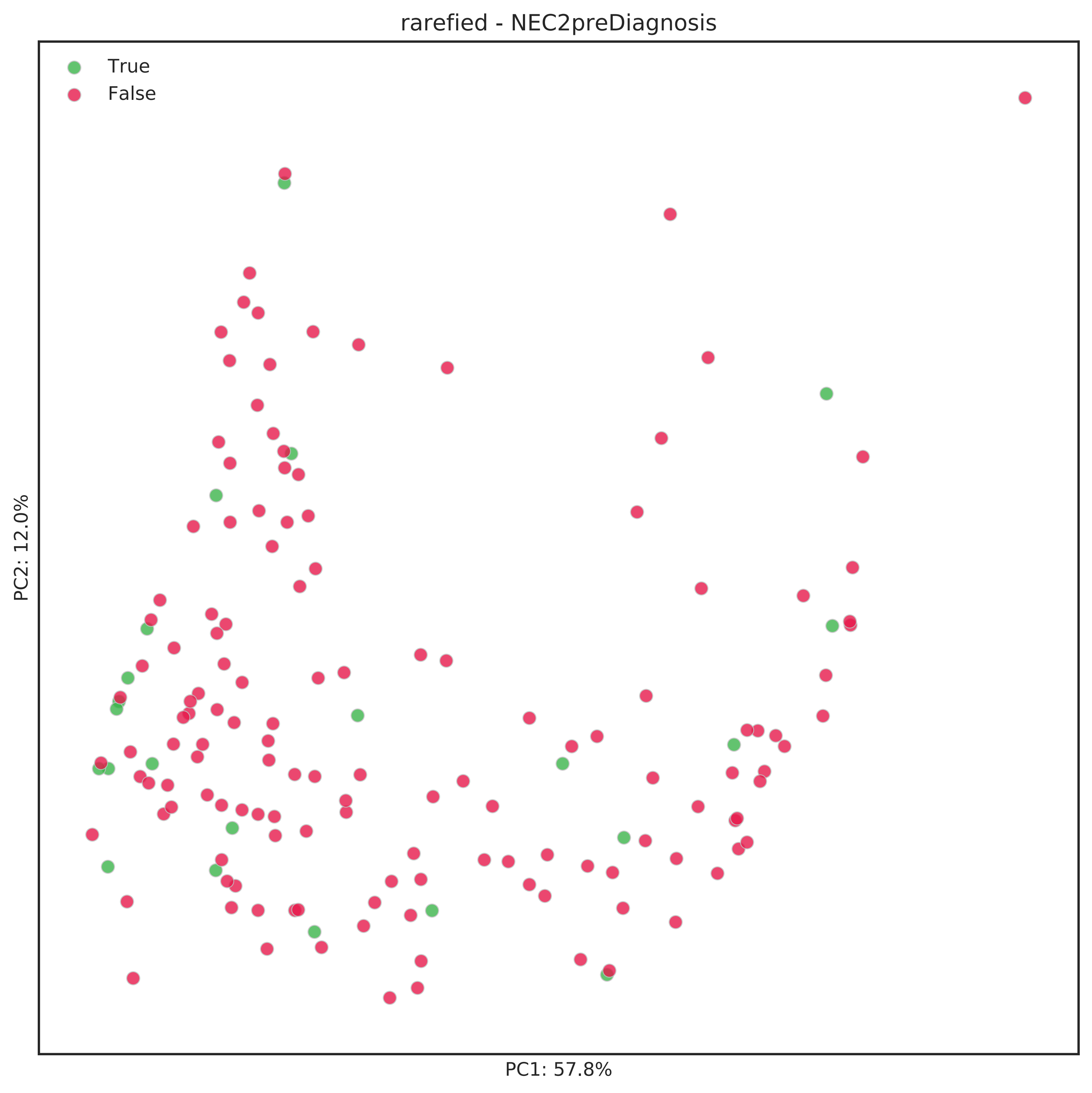
rarefied - NEC2preDiagnosis
True
False
PC2: 12.0%
PC1: 57.8%

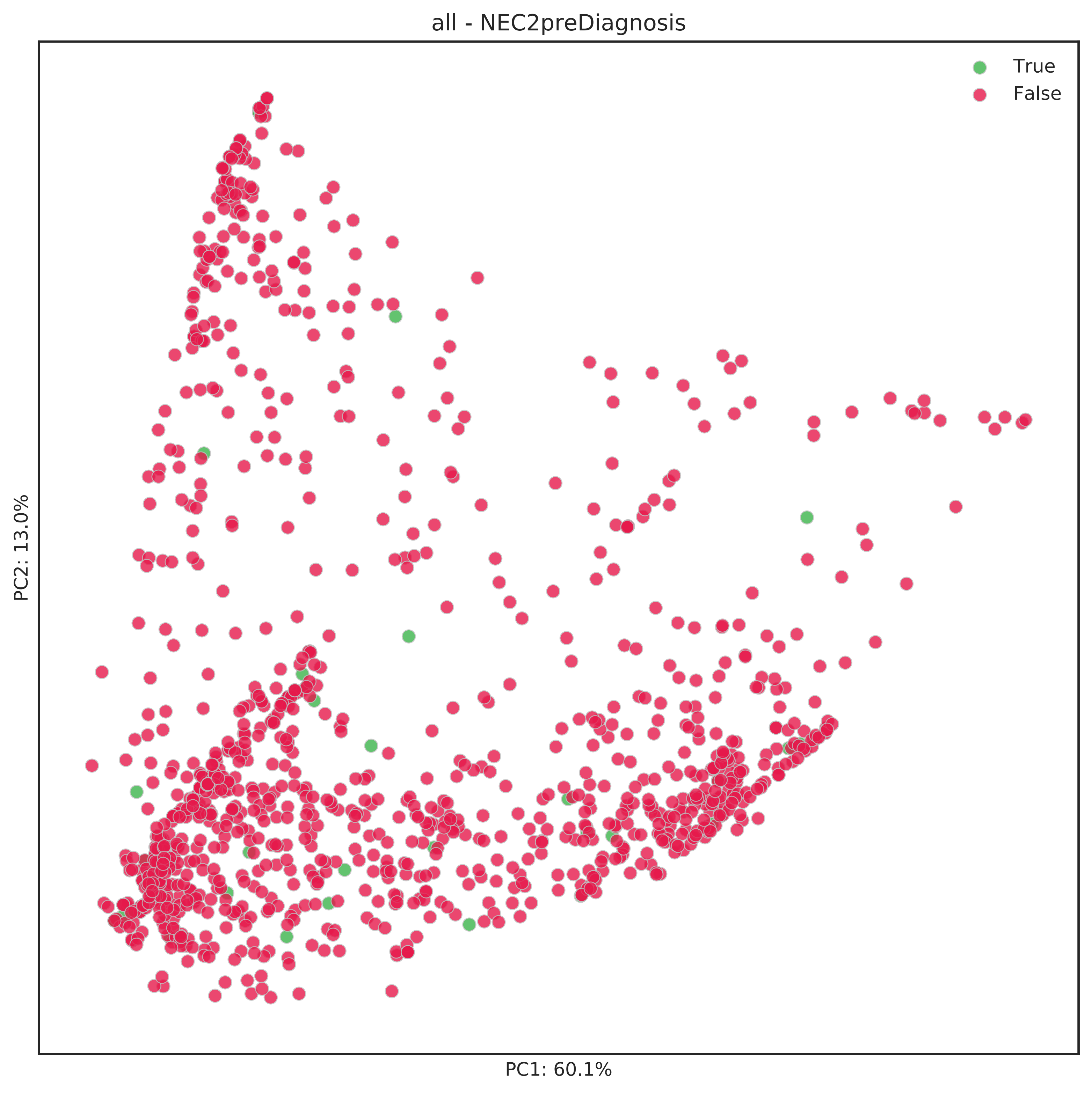
all - NEC2preDiagnosis
True
False
PC2: 13.0%
PC1: 60.1%

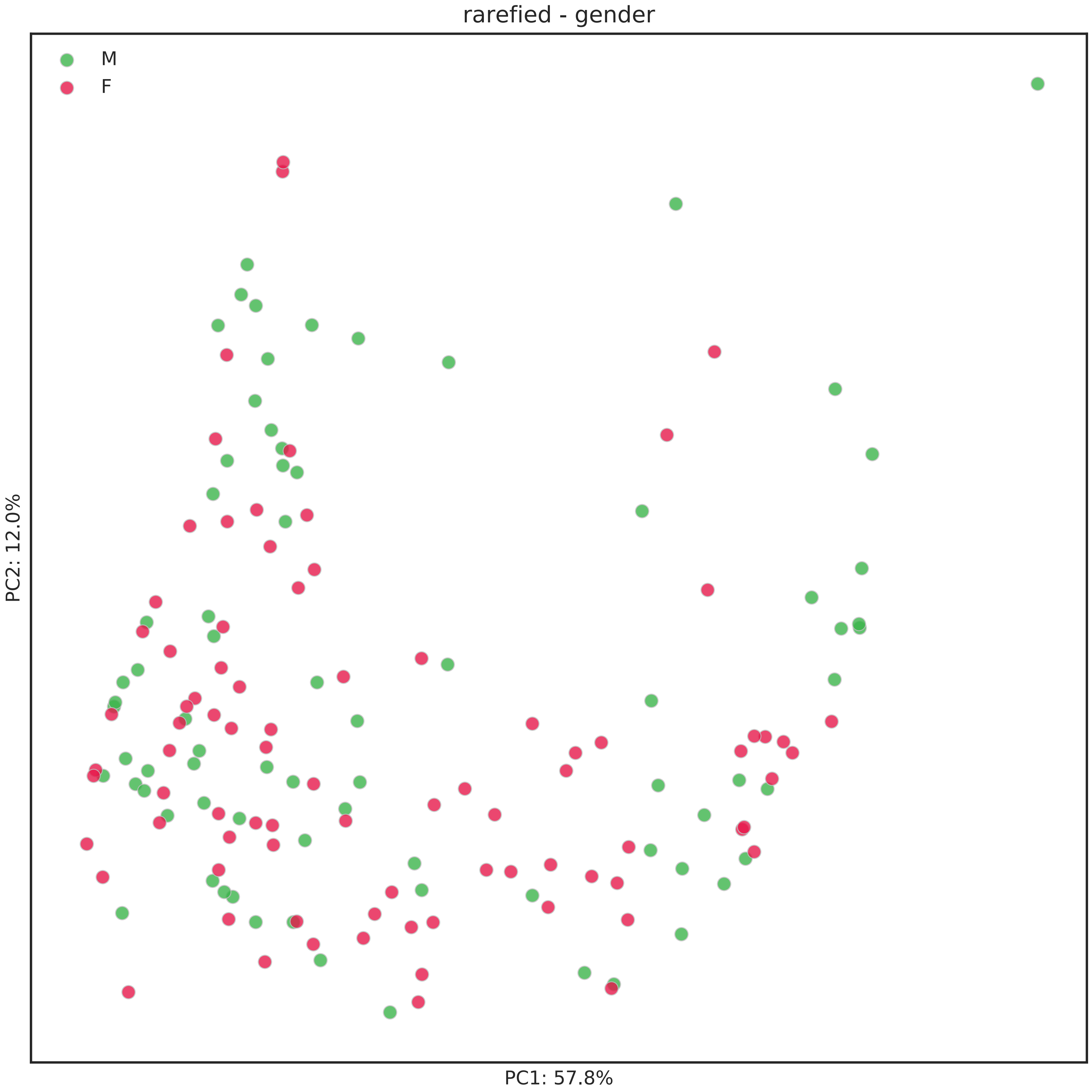
rarefied - gender
M
F
PC2: 12.0%
PC1: 57.8%

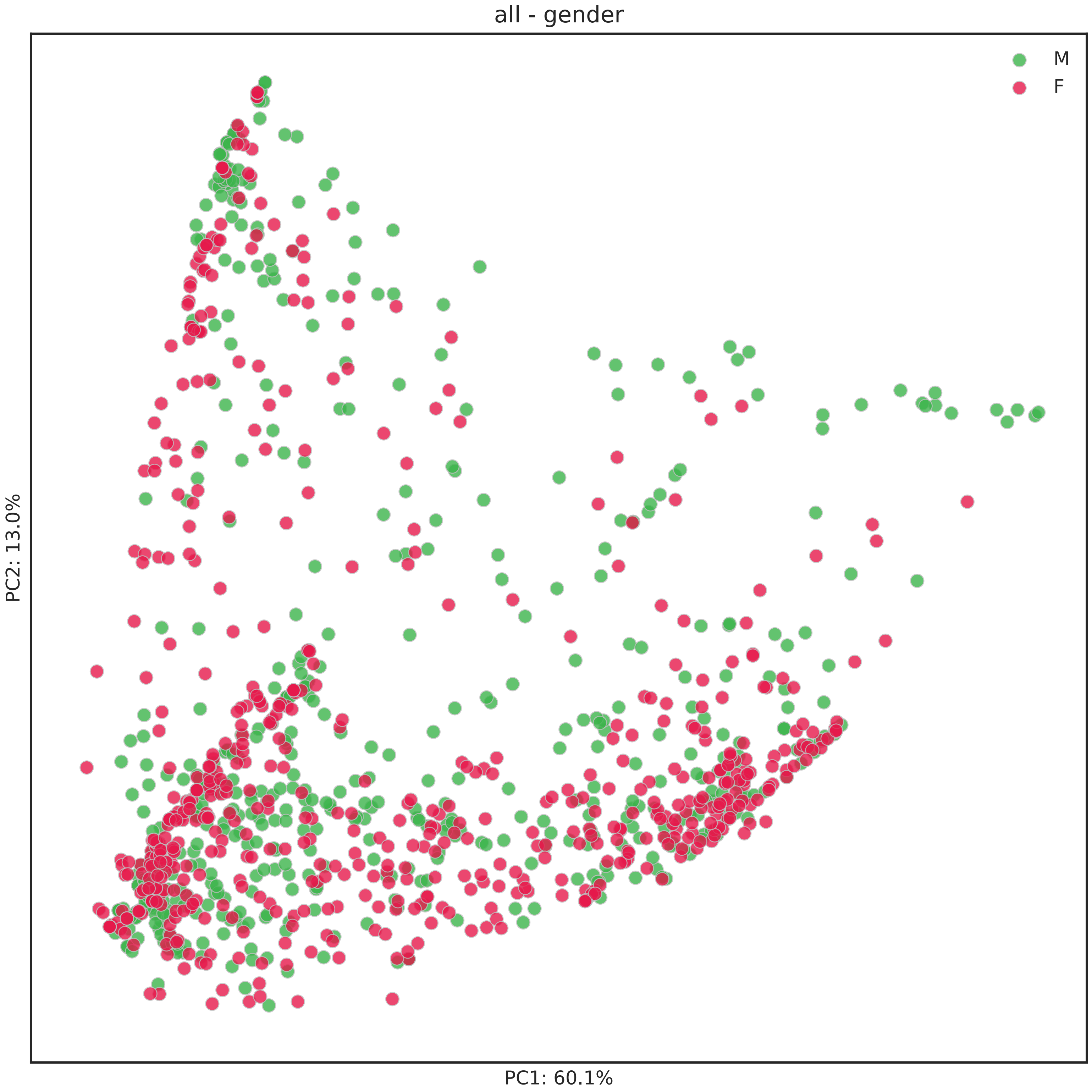
all - gender
M
F
PC2: 13.0%
PC1: 60.1%

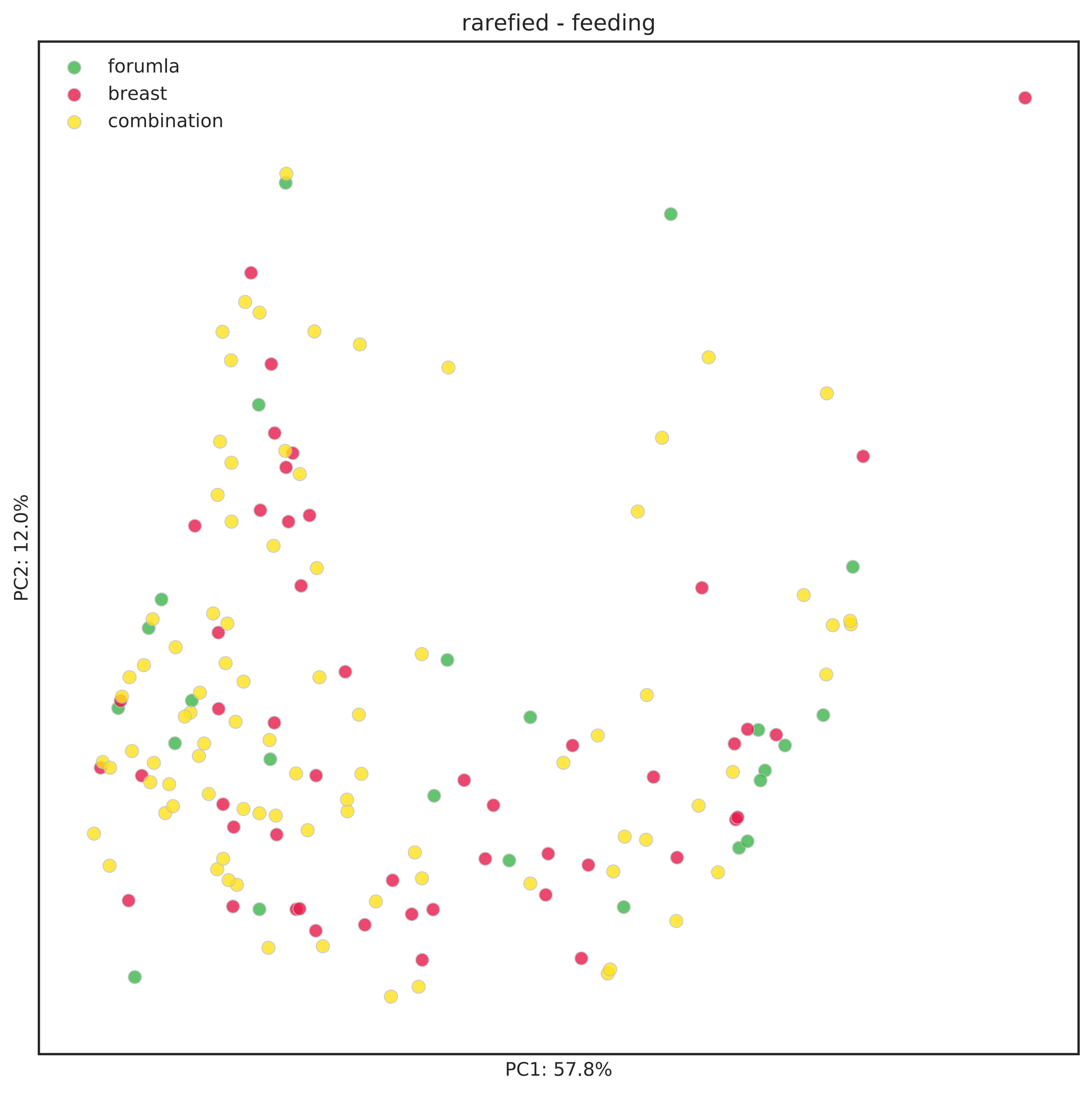
rarefied - feeding
forumla
breast
combination
PC2: 12.0%
PC1: 57.8%

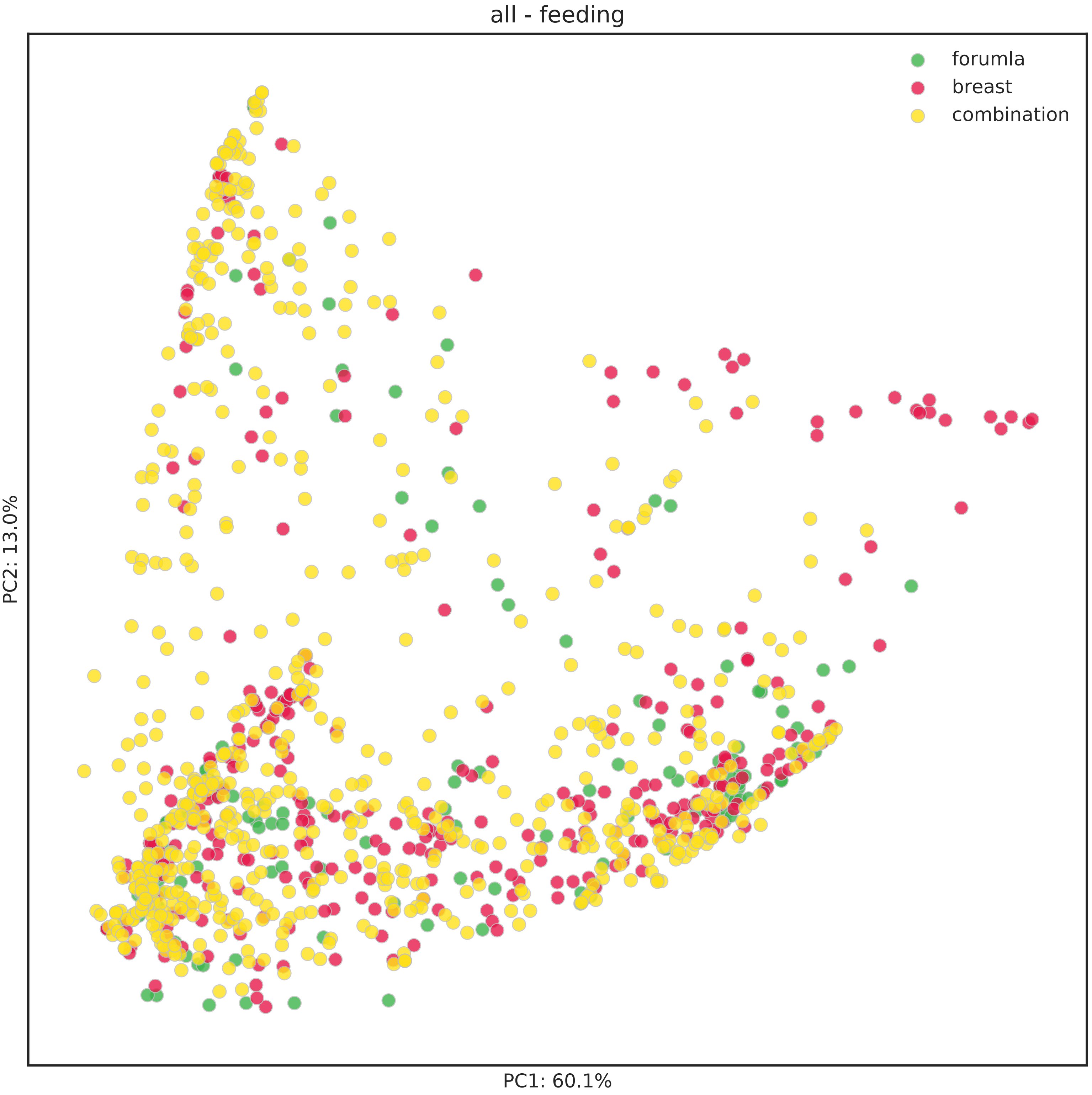
all - feeding
forumla
breast
combination
PC2: 13.0%
PC1: 60.1%

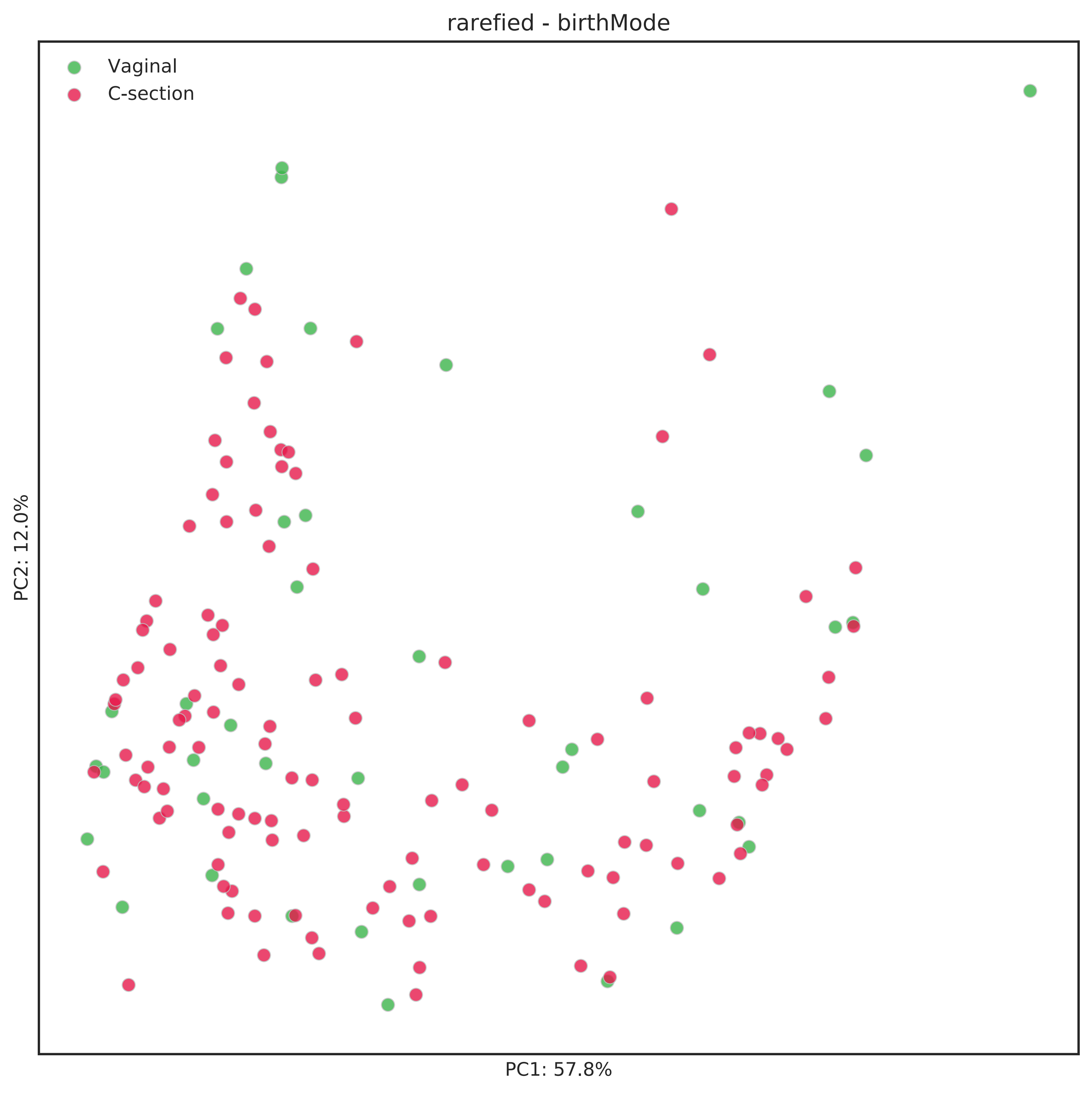
rarefied - birthMode
Vaginal
C-section
PC2: 12.0%
PC1: 57.8%

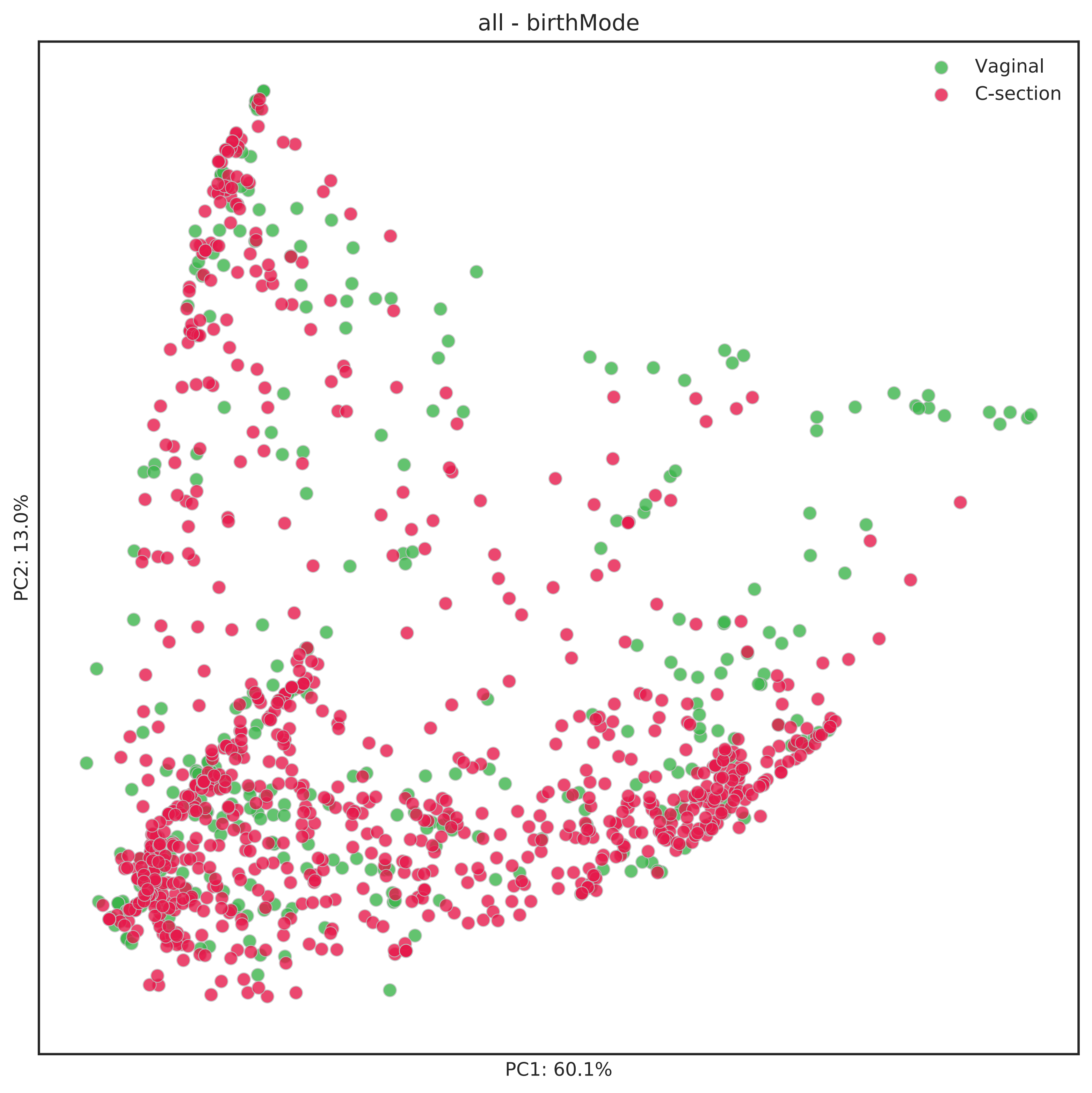
all - birthMode
Vaginal
C-section
PC1: 60.1%
PC2: 13.0%
